# Combining heterogeneous data sources for neuroimaging based diagnosis: re-weighting and selecting what is important

**DOI:** 10.1101/484311

**Authors:** Michele Donini, João M. Monteiro, Massimiliano Pontil, Tim Hahn, Andreas J. Fallgatter, John Shawe-Taylor, Janaina Mourão-Miranda, for the Alzheimer’s Disease Neuroimaging Initiative

## Abstract

Combining neuroimaging and clinical information for diagnosis, as for example behavioral tasks and genetics characteristics, is potentially beneficial but presents challenges in terms of finding the best data representation for the different sources of information. Their simple combination usually does not provide an improvement if compared with using the best source alone. In this paper, we proposed a framework based on a recent multiple kernel learning algorithm called EasyMKL and we investigated the benefits of this approach for diagnosing two different mental health diseases. The well known Alzheimer’s Disease Neuroimaging Initiative (ADNI) dataset tackling the Alzheimer Disease (AD) patients versus healthy controls classification task, and a second dataset tackling the task of classifying an heterogeneous group of depressed patients versus healthy controls. We used EasyMKL to combine a huge amount of basic kernels alongside a feature selection methodology, pursuing an optimal and sparse solution to facilitate interpretability. Our results show that the proposed approach, called EasyMKLFS, outperforms baselines (e.g. SVM and SimpleMKL), state-of-the-art random forests (RF) and feature selection (FS) methods.

## 1. Introduction

In this paper we study the problem of combining information from different data sources (e.g. imaging, clinical information) for diagnoses of psychiatric/neurological disorders. From a machine learning perspective, we have to solve a problem in a high dimensional space using only a small set of examples for training a predictive model. In the past few years, several papers investigated possible ways to combine heterogeneous data in neuroimaging-based diagnostic problems. Most of the previous approaches can handle only few different sources of information. The main goal of these approaches is to find an optimal combination of the sources in order to improve predictions, given different modalities of neuroimaging and other clinical information (as for example, demographic data or non-imaging biomarkers). In this context, Multiple Kernel Learning (MKL) provides an effective approach to combine different sources of information, considering each source of information as a kernel, and identifying which information is relevant for the diagnostic problem at hand [1, 2]. It is known that using multiple kernels instead of a single kernel can improve the classification performance (see e.g. [1] and references therein), and the goal of MKL is to find the correct trade-off among the different sources of information [1]. Moreover, MKL allows the extraction of information from the weights assigned to the kernels, highlighting the different importance of each different source. Therefore, applications of MKL to neuroimaging based diagnosis might help the discovery of biomarkers of neurological/psychiatric disorders.

### 1.1. Related Work

A number of recent studies have applied the MKL approach for multi-modal neuroimaging based diagnoses. Different MKL algorithms mainly differ on the type of kernels they use for each source (e.g. linear, Gaussian, polynomial) and on the way they estimate and combine the weights of the kernels. In general, most approaches impose some constraints on the norm^2^ of the weights (e.g. *ℓ_p_*-MKL [3]). In particular, the *ℓ*_1_-norm constraint imposes sparsity on the kernel combination therefore is able to select a subset of relevant kernels for the model (e.g. *ℓ*_1_-MKL [4]). The MKL framework is formally introduced in Section 2.

In [5] the authors exploit the standard *ℓ_p_*-MKL approach with p values rand-ing from 1 (sparse) to 2 (dense). They combine various sets of basic kernels (Gaussian, linear and polynomial) generated by selecting the top most relevant features (with the rank of the features determined by a t-test) extracted from Magnetic Resonance Imaging (MRI), Positron Emission Tomography (PET) images and clinical measurements. Their results show that this methodology outperforms the best kernel generated by exploiting the best unique source (MRI, PET or clinical measurements), suggesting that the combination of heterogeneous information with MKL is beneficial. Nevertheless, using a standard *ℓ_p_*-MKL approach imposes a limitation on the number of different basic kernels, due to the computational complexity and memory requirements of the *ℓ_p_*-MKL algorithms [1].

Another MKL approach able to combine different source of information is presented in [6], in which the authors tackle the problem of predicting the cognitive decline in older adults. In this case, the authors use the *ℓ_2_*-MKL with two Gaussian kernels, one for the MRI features and one for the clinical measurements. These kernels have two different hyper-parameters which were fixed by using a heuristic method. They claim that, by using only the MRI information or the clinical measurements alone, the kernels do not carry sufficient information to predict cognitive decline. On the other hand, using the kernel obtained by the combination of kernels extracted from both sources of information improves the performances significantly.

The problem of combining heterogeneous data for predicting Alzheimer’s disease has been handled also using the, so called, Multi-Kernel SVM. The idea is to use the standard SVM [7], with a pre-computed kernel that contains a weighted combination of the basic kernels. In this case, the combination is evaluated by exploiting a brute force search of the parameters (i.e. a grid search). In [8] and [9], the authors try to learn an optimal kernel combining three different kernels, each of which corresponds to a different sources of information (MRI, PET and clinical data), and the optimal (convex) combination of these kernels is determined via grid search. In [8] the authors propose, as first step of their methodology, a simple feature selection by using a t-test algorithm. In [9], the feature selection phase is improved by using a common subset of relevant features for related multiple clinical variables (i.e. Multi-Task learning approach [10]). In both studies, [8] and [9], the feature selection is applied before the generation of the kernels. Moreover, the brute force selection for the kernels weights, performed by using a grid search approach, is able to combine only few kernels and often finds a sub-optimal solution due to the manual selection of the search grid. In this sense, a MKL approach is more robust and theoretically grounded.

A recent paper by Xing Meng et al. [11] proposes a framework to predict clinical measures by using a multi-step approach. The authors combine three different neuroimaging modalities: resting-state functional Magnetic Resonance Image (fMRI), structural Magnetic Resonance Image (sMRI) and Diffusion Tensor Imaging (DTI). After a feature selection step within each of the single modalities, a selection of well-connected brain regions is performed. Their multi-modal fusion methodology consists of a simple concatenation of the selected features, ignoring the relative contribution of each modality. However, their approach does not include a weighting phase of the different modalities (in contrast with the MKL approach).

Other methodologies to combine different sources of information can be found in the literature [11–16]. One way is to exploit the Gaussian Processes for probabilistic classification (see e.g. [17]). For example, in [18], the authors combine five different modalities (i.e. segmentation of the brain in grey matter, white matter and cerebrospinal fluid, from T2 structural images plus the Fractional Anisotropy (FA) and Mean Diffusivity (MD) images, from the DTI sequence) to predict three Parkinsonian neurological disorders. Finally, in [19], the authors used Gaussian Processes to combine three different heterogeneous source of data: MRI, PET and the Apolipoprotein E (APOE) genotype, in order to predict conversion to Alzheimer’s in patients with mild cognitive impairment.In this family of methods, the Gaussian Process models have similarities with the MKL models, i.e. the goal is to find a kernel that combine prescribed kernels corresponding to each source of information plus a bias term. However, in these cases the models’ hyperparameters (kernels coefficients and bias terms) are selected using the Gaussian Process framework.

Another possible way to combine different sources of information is using RF-based methods [20, 21]. The framework used in these studies consists of several steps, where the RF methods are fundamental in order to obtain the final model as a combination of the different sources.

For example, the method proposed in [20] uses a RF model per modality in order to produce a similarity measure, one per source of information. Then, an approach to reduce of the number of features is applied and, in order to combine the data from different modalities, a selection of weights is performed by cross-validation. The output of this procedure is a weighted sum of the different measures of similarity that is equivalent to a combination of kernels, each one representing one modality.

As another example, the algorithm in [21] consists of a sequential exploitation of graph theory, recursive feature elimination (RFE) and RF. Graph theory is used to derived a set of features that are added to them the raw data. A RFE procedure is exploited in order to obtain a low dimensional set of features, one set per source of information. Then, one predictor per modality is generated by applying the RF to the selected features. Stacking all the resulting models (one per source of information) produces the final model.

In all previous studies outlined above, there is a limit on the maximum number of kernels that we are able to combine (or number of sources that we can consider) in the predictive model. In addition, feature selection (when performed) is applied before the generation of the final representation (i.e. the way how we describe the similarity among examples), thereby decreasing the connection between the final model and the selected features. These methods are not able to perform a fine-grained feature learning because they are heavily dependent on some priors (imposed by an expert), as for example the selection of which features are contained in a specific kernel.

### 1.2. Our contribution

In this paper, we proposed a MKL based approach that is able to re-weight and select the relevant information when combining heterogeneous data. This approach enables us to fragment the information from each data source into a very large family of kernels, learning the relevance of each fragmented information (kernel weights). Consequently, our method is very flexible and the final model is based on a kernel that uses a small amount of features, due to the feature selection performed as final step of our approach in synergy with the MKL methodology.

We start describing EasyMKL [22], a recent MKL algorithm, that can handle a large amount of kernels and we combine it in synergy with a new Feature Selection (FS) approach. Our aim is to evaluate and select the most relevant features from each data source. The proposed approach is applied to two different classification tasks. The first one considers the Alzheimer’s Disease Neuroimaging Initiative (ADNI) dataset to classify patients with Alzheimer’s disease vs. healthy controls combing structural MRI data and clinical assessments. Secondly, we tackle the task investigated in [23] where the goal is to classify depressed patients vs healthy controls by integrating fMRI data with additional clinical information. We compare our approach with SVM [7] as the baseline approach, as well as a state-of-the-art MKL approach (SimpleMKL [4]), two feature selection approaches: recursive feature elimination (RFE) [24] and t-test [25], and RF-based methods ([20, 21]).

In summary, the main contributions of this paper are two-fold. Firstly, we introduce a new methodology to combine a MKL approach using a huge number of basic kernels and a FS approach in order to improve the prediction performance, inherited from the previous preliminary work [26]. This new procedure, called EasyMKLFS, automatically selects and re-weights the relevant information obtaining sparse models. EasyMKLFS provides a new optimal kernel that can be used in every kernel machine (e.g. SVM) in order to generate a new classifier. Secondly, we demonstrate the performance of the proposed methodology using two classification tasks. When applied to the ADNI dataset the proposed approach was able to outperform the previous state-of-art methods and provide a solution with high level of interpretability (i.e. the identification of a small subset of features relevant for the predictive task); when applied to the depression dataset the proposed approach showed better performance than most approaches (a part from EasyMKL) with advantage of higher sparsity/interpretability.

The paper is organized as follows. In the first part of Section 2 we introduce the theory of MKL with an analysis of the most common MKL methods. Then, the original EasyMKL is presented, followed by the connection between feature learning and MKL. The proposed method is described in the last part of section Section 2.4. Section 3 shows the main information about the datasets, the methods, the validation procedure for the hyper-parameters and the details concerning the performed experiments. Section 4 describes the datasets used in this study, the methods used as comparisons against EasyMKLFS, the validation procedure, and the experimental designs. The results are presented in Section 4 for both datasets, followed by a discussion in Section 5.

## 2. Theory

In the next sections, we will introduce the classical MKL framework and a recent MKL algorithm called EasyMKL. Firstly, we introduce the notation used in this paper.

Considering the classification task, we define the set of the training examples as 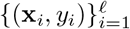 with x_*i*_ in 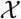 and *y_i_* with values +1 or −1. In our case, it is possible to consider the generic set 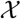 equal to 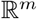, with a very large number of features *m*. Then, 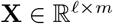 denotes the matrix where examples are arranged in rows. The *i^th^* example is represented by the *i^th^* row of **X**, namely **X**[*i*,:] and the *r^th^* features by the *r^th^* column of **X**, namely **X**[:,*r*].

Specifically, in our cases, the number of examples *ℓ* refers to the number of different subjects that are considered in the study.

### 2.1. Multiple Kernel Learning (MKL)

MKL [1, 27] is one of the most popular paradigms used to highlight which information is important, from a pool of *a priori* fixed sources. The goal of MKL is to find a new optimal kernel in order to solve a specific task. Its effectiveness has been already demonstrated in several real world applications [28, 29]. A kernel **K** generated by these techniques is a combination of a prescribed set of *R* basic kernels **K**_1_,…, **K**_*R*_ in the form:

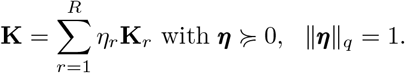

The value *q* defines the used norm and is typically fixed to 1 or 2. When *q* is fixed to 1, we are interested in a sparse selection of the kernels. However, if *q* equals 2, then the model will be dense (with respect to the selected kernels). It is important to highlight how the value *η_r_* represents the weight assigned to the specific *r^th^* source of information.

Using this formulation, we are studying the family of weighted sums of kernels. It is well known that the sum of two kernels is equivalent to the concatenation of the features contained in both the feature spaces [30]. Extending the same idea, the weighted sum of a list of basic kernels can be seen as a weighted concatenation of all the features contained in all the feature spaces (where the weights are the square roots of the learned weights *η_r_*).

Theoretically, MKL algorithms are supported by several results that bound the *estimation error* (i.e. the difference between the true error and the empirical margin error) [31–38].

#### 2.1.1. An overview of the MKL approaches

Existing MKL approaches can be divided in two main categories. In the first category, *Fixed or Heuristic*, some fixed rule is applied to obtain the kernel combination. These approaches scale well with the number of basic kernels, but their effectiveness critically depend on the domain at hand. They use a parameterized combination function and find the parameters of this function (i.e. the weights of the kernels) generally by looking at some measure obtained from each kernel separately, often giving a suboptimal solution (since no information sharing among the kernels is exploited).

Alternatively, *Optimization based* approaches learn the combination parameters (i.e. the kernels’ weights) by solving a single optimization problem directly integrated in the learning machine (e.g. exploiting the generalization error of the algorithm) or formulated as a different model, as for example by alignment, or other kernel similarity maximization [4, 27, 39].

In the *Fixed or Heuristic* family there are some very simple (but effective) solutions. In fact, in some applications, the average method (that equal to the sum of the kernels [40]) can give better results than the combination of multiple SVMs each trained with one of these kernels [41]. Another solution, can be the element-wise product of the kernel matrices contained in the family of basic kernels [42].

The second family of MKL algorithms is defined exploiting an optimization problem. Unexpectedly, finding a good kernel by solving an optimization problem turned out to be a very challenging task, e.g. trying to obtain better performance than the simple average of the weak kernels is not an easy task^3^. Moreover, *Optimization based* MKL algorithms have a high computational complexity, for example using semidefinite programming or quadratically constrained Quadratic Programming (QP). Some of the most used MKL algorithms are summarized in Table 1 with their computational complexities.

**Table 1:** Frequently used MKL *Optimization based* methods.

|  | Learner | Time Complexity | Reference |
| --- | --- | --- | --- |
| SimpleMKL | SVM | Grad.+ QP | Rakotomamonjy et al. [4] |
| GMKL | SVM | Grad.+ QP | Varma and Babu [39] |
| GLMKL | SVM | Analytical + QP | Kloft et al. [3] |
| LMKL | SVM | Grad.+ QP | Gönen and Alpaydin [43] |
| NLMKL | KRR | Grad.+ Matrix Inversion | Cortes et al. [44] |

### 2.2. EasyMKL

EasyMKL [22] is a recent MKL algorithm able to combine sets of basic kernels by solving a simple quadratic optimization problem. Besides its proved empirical effectiveness, a clear advantage of EasyMKL compared to other MKL methods is its high scalability with respect to the number of kernels to be combined. Specifically, its computational complexity is constant in memory and linear in time.

This remarkable efficiency hardly depends on the particular input required by EasyMKL. In fact, instead of requiring all the single kernel matrices (i.e. one per source of information), EasyMKL needs only the (trace normalized) average of them. See Section Appendix A (in the Appendix) for a technical description of EasyMKL^4^.

### 2.3. Feature Learning using MKL

In the last years, the importance of combining a large amount of kernels to learn an optimal representation became clear [22]. As presented in the previous section, new methods can combine thousands of kernels with acceptable computational complexity. This approach contrasts with the previous idea that kernel learning is shallow in general, and often based on some prior knowledge of which specific features are more effective. Standard MKL algorithms typically cope with a small number of strong kernels, usually less than 100, and try to combine them (each kernel representing a different source of information of the same problem). In this case, the kernels are individually well designed by experts and their optimal combination hardly leads to a significant improvement of the performance with respect to, for example, a simple averaging combination. A new point of view is instead pursued by EasyMKL, where the MKL paradigm can be exploited to combine a very large amount of basic kernels, aiming at boosting their combined accuracy in a way similar to feature weighting [2]. Moreover, theoretical results prove that the combination of a large number of kernels using the MKL paradigms is able to add only a small penalty in the *generalization error*, as presented in [31, 33–35].

In this sense, we are able to take a set of linear kernels that are evaluated over a single feature, making the connection between MKL and feature learning clear. The single kernel weight is, in fact, the weight of the feature. Using this framework, we can weight the information contained into a set of features, evaluated in different ways (i.e. using different kernels that can consider different subsets of features).

### 2.4. EasyMKL and Feature Selection

In this section, we present our approach to combine MKL (as a feature learning approach) and feature selection (FS). We start from EasyMKL with a large family of linear single-feature kernels as basic kernels. We decided to exploit linear kernels because they do not need hyper-parameter selection. Dealing with small datasets, this is a serious advantage. Moreover, in our single-feature context, using other families of kernels (e.g. RBF or polynomial kernels) has not impact on the final performances^5^. Due to the particular definition of this algorithm, we are able to combine efficiently millions of kernels. As presented in Section 2.2 and in Appendix A, given the kernel generated by the average of the trace normalized basic kernels

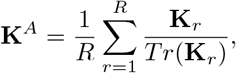

EasyMKL produces a list of weights 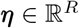, one weight per kernel.

Fixing a threshold *ρ* > 0, it is possible to remove all the kernels with a weight less or equal to *ρ*, considering them not sufficiently informative for our classification task. In this way we are able to inject sparsity in our final model. All the single-feature kernels **K**_*r*_ with a weight *η_r_* > *ρ* are weighted and summed obtaining a new kernel

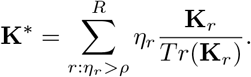

Algorithm 1 summarizes our approach, called EasyMKLFS. It is important to note that if *ρ* = 0 we are performing the standard MKL approach over *R* basic kernels.

The same procedure cannot be easily exploited with the standard MKL algorithms, due to the large amount of memory required to combine a large family of kernels (see Table 1). In this sense, EasyMKL becomes fundamental in order to efficiently achieve our goal. In line 8 of Algorithm 1, the amount of memory required by the storage of the kernels is independent with respect to the number of combined kernels *R* (and the computational complexity is linear in time).

#### Algorithm 1 **EasyMKLFS**: feature selection and weighting by using EasyMKL. 𝕆_ℓ,ℓ_ is the zero-matrix in ℝ^ℓ×ℓ^.

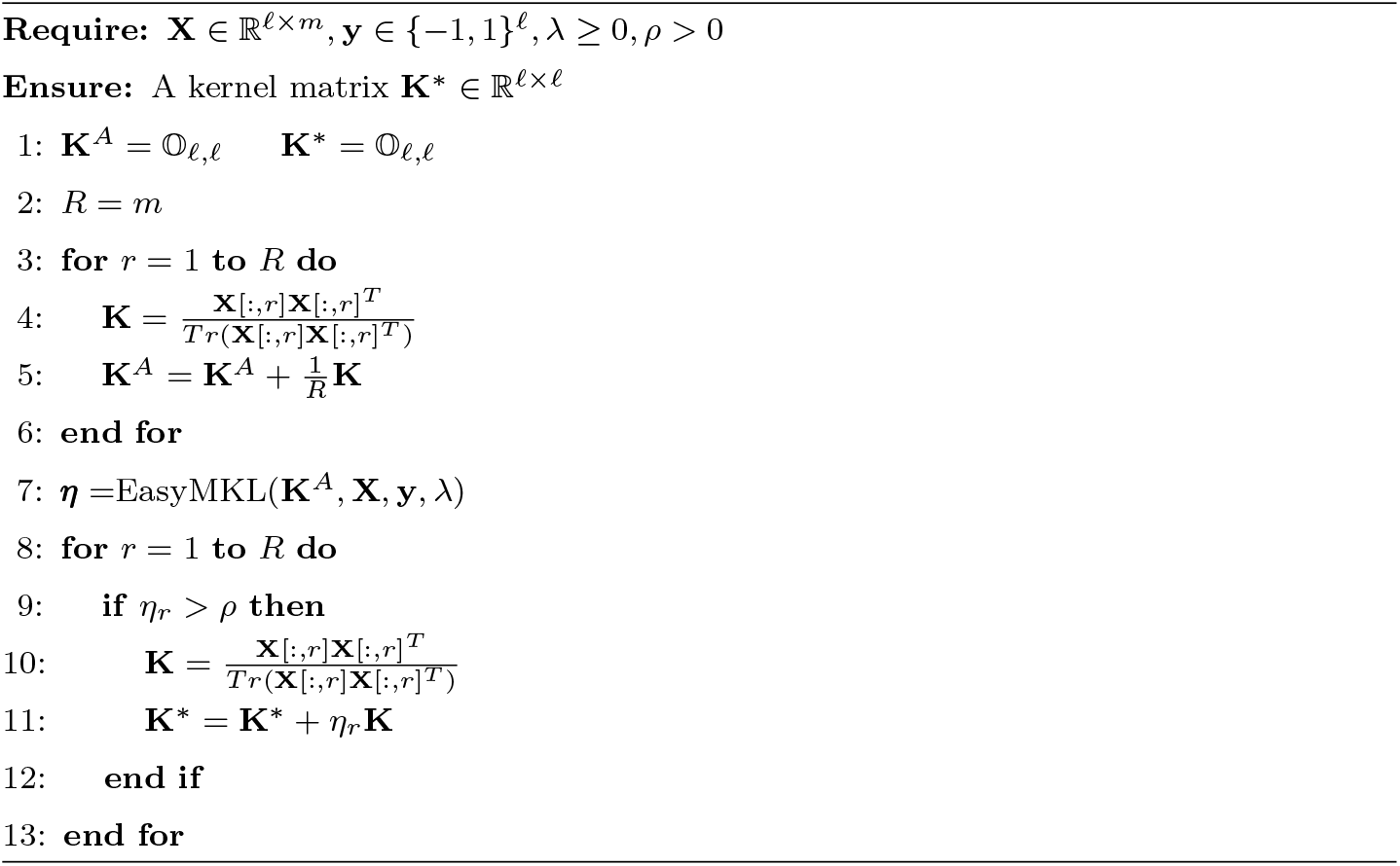

## 3. Materials and Methods

### 3.1. Datasets

In this section, we present a description of the two considered datasets, i.e. ADNI and Depression. The first dataset consists of structural Magnetic Resonance Imaging (sMRI), clinical and genetic information for each participant. The second dataset consists of functional MRI (fMRI) and clinical information for each participant.

#### 3.1.1. ADNI

This case study concerns the problem of classifying patients with possible Alzheimer’s disease combining sMRI images and other genetical/clinical or demographic information. Alzheimer’s disease (AD) is a neurodegenerative disorder that accounts for most cases of dementia.

In 2003, the ADNI was started as a public-private partnership by Principal Investigator Michael W. Weiner, MD. The primary goal of ADNI has been to test whether serial Magnetic Resonance Imaging (MRI), Positron Emission Tomography (PET), other biological markers, and clinical and neuropsychological assessment can be combined to measure the progression of mild cognitive impairment (MCI) and early Alzheimers disease (AD).

Here, we use sMRI and clinical information from a subset of 227 individual from the ADNI dataset. The following pre-processing steps were applied to sMRI of the selected individuals. The T1 weighted MRI scans were segmented using SPM12 into gray matter, white matter and Cerebral Spinal Fluid (CSF). The grey matter probability maps were normalised using Dartel, converted to MNI space with voxel size of 2mm × 2mm × 2mm and smoothed with a Gaussian filter with 2 mm FWHM. A mask was then generated, to select voxels which had an average probability of being grey matter equal or higher than 10% for the whole dataset. This resulted in 168130 voxels per subject being used.

Finally, from the non-imaging information contained in ADNI, we extracted 35 different clinical information, including age and gender of the patient, the presence of APOE4 allele, items of the *Mini-mental State Exam* (MMSE) [45], education level, *Clinical Demential Rating, AD Assessment Schedule 11* and 13, Rey Auditory Verbal Learning Test and *Functional Assessment Questionnaire* [46] (see Appendix A, Table B.12 for the details).

For up-to-date information about the ADNI, see www.adni-info.org.

#### 3.1.2. Depression

The task in this challenging dataset [23] is to classify depressed patients versus healthy controls by integrating fMRI data and other clinical measurements.

A total of 30 psychiatric in-patients from the University Hospital of Psychiatry, Psychosomatics and Psychotherapy (Wuerzburg, Germany) diagnosed with recurrent depressive disorder, depressive episodes, or bipolar affective disorder based on the consensus of two trained psychiatrists according to ICD-10 criteria (DSM-IV codes 296.xx) participated in this study. Accordingly, self report scores in the German version of the Beck Depression Inventory (second edition) ranged from 2 to 42 (mean standard deviation score, 19.0 [9.4]). Exclusion criteria were age below 18 or above 60 years, co-morbidity with other currently present Axis I disorders, mental retardation or mood disorder secondary to substance abuse, medical conditions as well as severe somatic or neurological diseases. Patients suffering from bipolar affective disorder were in a depressed phase or recovering from a recent one with none showing manic symptoms. All patients were taking standard antidepressant medications, consisting of selective serotonin reuptake inhibitors, tricyclic antidepressants, tetracyclic antidepressants, or serotonin and noradrenalin selective reuptake inhibitors. Thirty comparison subjects from a pool of 94 participants previously recruited by advertisement from the local community were selected to match the patient group in regard to gender, age, smoking, and handedness using the optimal matching algorithm implemented in the MatchIt package for R http://www.r-project.org [47]. In order to exclude potential Axis I disorders, the German version of the Structured Clinical Interview for DSM-IV (SCID; 35) Screening Questionnaire was conducted. Additionally, none of the control subjects showed pathological Beck Depression Inventory (BDI II) scores (mean = 4.3, SD = 4.6).

From all 60 participants, written informed consent was obtained after complete description of the study to the subjects. The study was approved by the Ethics Committee of the University of Wuerzburg, and all procedures involved were in accordance with the latest version (fifth revision) of the Declaration of Helsinki.

The fMRI task consisted of passively viewing four types of emotional faces. Anxious, Happy, Neutral and Sad facial expressions were used in a blocked design, with each block containing pictures of faces from 8 individuals obtained from the Karolinska Directed Emotional Faces database: http://www.emotionlab.se/resources/kdefdatabase. Every block was randomly repeated 4 times. Subjects were instructed to attend to the faces and empathise with the emotional expression. Images acquisition details can be found in previous publications using this dataset [23].

The images were preprocessed using the Statistical Parametric Mapping software (SPM5, Wellcome Department of Cognitive Neurology, U.K.). Slice-timing correction was applied, images were realigned, spatially normalised and smoothed using an 8 mm FWHM Gaussian isotropic kernel. For each participant, a General Linear Model (GLM) was applied in which each emotion was modeled by the convolution of the blocks with the hemodynamic response function. The contrast images corresponding to each emotion were used for the classification models. More specifically, for each subject we combined four different contrast images, corresponding to the brain activations to the four different emotional faces: Anxious, Happy, Neutral and Sad.

From the non-imaging information contained in the Depression dataset, we generated a list of 48 different clinical and demographic variables, including age, gender and several results from psychological tests as *Karolinska Directed Emotional Faces* [48] test, the *Sensitivity to Punishment/Reward Questionnaire* [49], tests of processing speed (approx. IQ) [50], *Montgomery-Asberg depression rating scale* [51], *Self-report questionnaire of depression severity* [52], *Positive-Negative Affect Schedule* [53] and *State-Trait inventory* [54] (see Appendix A, Table B.13 for the complete list).

### 3.2. Experimental settings

We combine features derived from the images (each voxel is considered as a single feature) with sets of selected clinical and demographic features. In the following we will refer to (linear single-feature) basic kernels or directly to features without distinction.

In our experiments, we consider different subsets and different fragmentations of the whole information contained in the datasets. The considered linear kernels (or features) are divided in 7 different sets:

- **I** represents all image features in one single linear kernel (in case of the fMRI dataset which contains 4 images it corresponds to concatenate all the features in only one kernel).
- **C** represents the whole clinical/demographic information in one single linear kernel.
- **I+C** is the kernel containing all the voxels and all the clinical/demographic features, which corresponds to the simplest way of combining (or concatenating) the different sources.
- **I & C** is the grouping of information with one group for each imaging information (MRI or fMRI) each one containing all the voxels and one group for the clinical/demographic information. This way of grouping the data is exploited in the context of RF methods, in order to maintain a feasible computational complexity.
- **I** & 𝒞 is the family of basic kernels that contains a single linear kernel for each whole image (i.e. one kernel per image) plus one kernel for each clinical/demographic feature. In this case, we are able to tune the importance of the single clinical feature, and make the correct trade-off between clinical information and image information.
- 𝒱 is the family of basic kernels (or basic features) that contains one kernel for each voxel. Each single voxel can be weighted or selected, pointing out the relevant voxels of the MR images.
- 𝒱 & 𝒞 is the family of basic kernels (or basic features) that contains one kernel for each voxel plus one kernel for each clinical feature. This is the most flexible model which is able to point out the relevant voxels and clinical/demographic features.

Our new methodology exploits the 𝒱 & 𝒞 set and it can be divided in three principal steps. The first step is the extraction of the features and their vectorization. Then, as a second step, we apply our algorithm (EasyMKLFS) to weight and select the features. Finally, we are able to generate a sparse (linear) model by using the obtained kernel in a classifier (e.g. SVM). The idea behind our methodology is summarized in Figure 1. Specifically, in the present work we used the SVM as a classifier as it is a machine learning algorithm that performs very well in many different type of problems.

**Figure 1:**
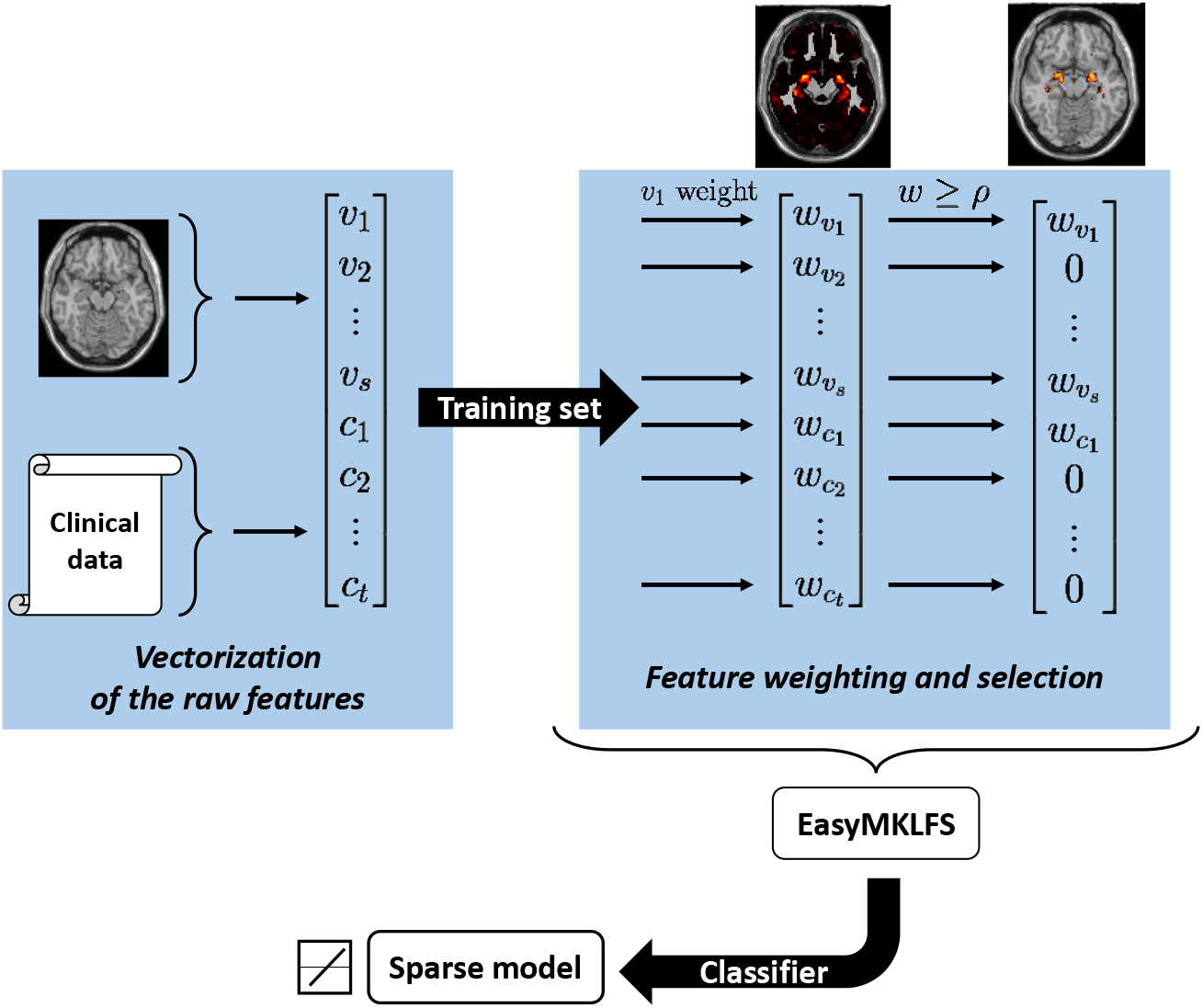
Our framework with the three principal steps: (1) extraction of the raw features (from MRIs, i.e. *v*_1_,…,*v_s_*, and from clinical data, i.e. *c*_1_,…,*c_t_*); (2) evaluation of the important information by using EasyMKLFS for feature weighting and selection; (3) generation of the final sparse model.

### 3.3. Comparison with other methods

We performed a balanced accuracy comparison (i.e., the average between sensitivity and specificity) considering 6 different families of methods:

- *Baseline:* Linear SVM [7], using the linear kernels generated using the whole images (**I**), clinical information (**C**) or both (**I + C**). It is used as baseline to understand the challenge of the classification tasks.
- *FS:* the second family of approaches is comprised of two feature selection (FS) methods. We applied these algorithms considering each voxel of the images as a single feature (𝒱) or adding both one feature per voxel and one feature for each clinical information (𝒱 & 𝒞). The first method is the SVM RFE [24], which corresponds to the standard recursive feature elimination approach. RFE considers the importance of individual features in the context of all the other features, and it has the ability to eliminate redundancy, and improves the generalization accuracy [55]. The second one is the SVM t-test, a heuristic method that exploits a statistical test for evaluating the importance of the features. The selected features are then used in a SVM. FS method is univariate and it is not able to take into account the interactions between features [25].
- *RF:* the third comparison is with respect to the RF-based approaches. The RF methods select the relevant features, in each modality, independently with respect to the other sources of information. In this sense, we consider RF exploiting the **I** & **C** as segmentation of the sources of information in order to highlight the differences compared to the other presented methodologies. We implement two methods, namely Gray [20] and Pustina [21], where the RF algorithms are the key in order to find the best representation of the single source of information. These methods are not kernel-based methods, and are composed by a pipeline of different algorithms. We tried to make the comparison as fair as possible, but we are aware that the same authors in [20] highlighted that a direct comparison with other existing methods is hard to perform due to problems such as the inclusion of different subjects and modalities, as well as the use of different methods for feature extraction and cross-validation. Moreover, we highlight that the computational complexity of these methods is significantly higher than the others. For this reason, they are not able to handle a larger number of different sources of information.
- *MKL:* the fourth comparison is against the standard MKL methodology. Firstly, we used SimpleMKL [4], a well known MKL iterative algorithm that implements a linear approach based on a sparse combination of the kernels. Secondly, we used EasyMKL, a recent MKL algorithm presented in Section 2.2 and Appendix A. We provided to these algorithms a family of basic kernels composed by one kernel per image and one kernel per clinical information **I** & 𝒞 (i.e. a small family of basic kernels).
- *FW:* in this group we applied a different point of view for the MKL [22]. In this new context, we consider MKL as a feature weighting algorithm and we provide to EasyMKL a single kernel for each feature (voxels and clinical information, i.e. 𝒱 & 𝒞). We are not able to compare EasyMKL with SimpleMKL in this setting, because of the computational and memory requirement of this algorithm.
- *FWS:* the last comparison is our EasyMKLFS, which consists in a combination of MKL with FW and FS, as described in Section 2.4. We tested our method with one kernel per voxel (𝒱), and one kernel per voxel plus one kernel per clinical information (𝒱 & 𝒞) as basic kernels.

The kernels, generated by *MKL, FW* and *FWS* methods, are plugged into a standard SVM. In this way, we are able to compare the quality of different kernels avoiding the possible noise given by different classifiers. As highlighted before, the RF-methods are based on a different classifier. In the following, we tried to maintain the comparisons as fair as possible.

It is important to highlight that our approach, similarly to the other approaches used for comparison, have the following two main assumptions: (i) there are features in the data that are able to distinguish between two groups, despite of their within-group heterogeneity. (ii) different sources of information might carry complementary information for the classification task and, consequently, combining them can be advantageous.

For both datasets, we used the Wilcoxon signed-rank test [56] to compare the proposed algorithm (EsasyMKLFS) with the other methods. More specifically, we tested whether the proposed algorithm provided statistically significant different predictions with respect to the other methods. We used the Bonferroni correction to account for multiple comparisons, therefore the p-value threshold for rejecting the null hypothesis that two classifiers are not different was 0.05 divided by the number of comparisons (i.e. 8 for both datasets).

#### 3.3.1. Validation

All the experiments are performed using an average of 5 repetitions of a classic nested 10-fold cross-validation. We fixed the same distribution of the age of the patients among all the subsets.

The validation of the hyper-parameters has been performed in the family of *C* ∈ {0.1,1, 5, 25} for the SVM parameter, 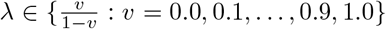 for the EasyMKL parameter, 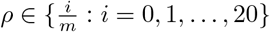 (where *m* is the number of the features) for the EasyMKLFS parameter. We fixed the percentage of dropped features at each step of the feature selection approaches (RFE and t-test) equal to the 5% (using higher percentages deteriorates the results).

Specifically, we reported the average of 5 repetitions of the following procedure:

- The dataset is divided in 10 folds **f**_1_,…,*f*_10_ respecting the distribution of the labels and the age of the patients, where **f**_*i*_ contains the list of indexes of the examples in the *i*-th fold;
- One fold **f**_*j*_ is selected as test set;
- The remaining nine out of ten folds 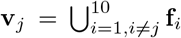 are then used as validation set for the choice of the hyper-parameters. In particular, another 10-fold cross validation over **v**_*j*_ is performed (i.e., nested 10-fold cross-validation);
- The set **v**_*j*_ is selected as training set to generate a model (using the validated hyper-parameters);
- The test fold **f**_*j*_ is used as test set to evaluate the performance of the model;
- The collected results are the averages (with standard deviations) obtained repeating the steps above over all the 10 possible test sets **f**_*j*_, for each *j* in {1,…, 10}.

#### 3.3.2. Clinical information settings

We considered two different experimental settings. Firstly, we removed the clinical information which are highly correlated with the labels. Note that, in both cases, dementia and depression, the diagnosis or labels are derived from clinical measures due to the lack of biomarkers, therefore by excluding clinical information highly correlated with the labels we are basically avoiding circularity or double dipping in the analysis. We performed a t-test between each individual feature and the corresponding label, and then excluded the ones that were statistically correlated with the labels by using *p* < 0.01 with false discovery rate (FDR) correction for multiple comparisons. FDR [57] is a powerful method for correcting for multiple comparisons that provides strong control of the family-wise error rate (i.e., the probability that one or more null hypotheses are mistakenly rejected).

The remaining clinical information after this selection are 25 for the ADNI dataset and 44 for Depression dataset. The idea is to show that the improvement of the results is not due to the use of clinical variables which are directly used by experts to assign the patient labels.

In the second set of experiments, we used all the clinical variables available. The results of these experiments can be found in the supplementary material, as a sanity check of our datasets and methodologies. A large increase of accuracy is obtained from this second experiment. However, these results can be considered over optimistic, as the clinical features are highly correlated with the labels.

### 3.4. Weight Maps Summarization

In the present work we used a method described in [58] to rank the regions that contribute most to the predictive model according to the Automated Anatomical Labeling (AAL) Atlas [59]. More specifically, the regions were ranked based on the average of the absolute weight value within them.

Therefore, regions which contain weights with a large absolute value, and/or contain several weights with values different from zero, will be ranked higher.

## 4. Results

In this section, the results are summarized for both the datasets. When it is reasonable, we firstly compare all the presented methods considering only the image or clinical features. Secondly, we compare different methods to combine heterogeneous data, i.e. images and clinical/demographic information.

### 4.1. ADNI

In this section we present the results obtained using the ADNI dataset. The results are presented for the previously described methods: Baseline (i.e. linear SVM), Feature Selection (FS), Random Forests methods (RF), Multiple Kernel Learning (MKL), Feature Weighting by using MKL (FW) and the proposed method Feature Weighting and Selection (FWS). In Table 2 the results obtained by exploiting only one source of information are reported, i.e. clinical information or features derived from structural MRI. It is possible to see that the SVM algorithm with only the clinical information is not able to generate an effective predictive model. Due to the small amount of clinical features (with respect to the examples), using FS or FW would not be effective, therefore, this comparison will not be presented. Concerning the MR images, there is a small increase in balanced accuracy when using either feature selection, feature weighting, or both.

**Table 2:** ADNI Dataset: comparisons of 5 repetitions of a nested 10-fold cross-validation balanced accuracy using the clinical information selected by a FDR procedure. The results are divided in 4 families: Baseline, Feature Selection (FS), Feature Weighting by using MKL (FW) and our method in Feature Weighting and Selection (FWS). R corresponds to the number of kernels used.

|  | Algorithm | Kernels | R | Bal. Acc. % |
| --- | --- | --- | --- | --- |
| Baseline | SVM | $\mathbf{C}$ | 1 | $52.12 \pm 8.26$ |
| | SVM | $\mathbf{I}$ | 1 | $84.08 \pm 6.94$ |
| FS | SVM RFE | $\mathcal{V}$ | — | $86.34 \pm 6.93$ |
| | SVM t-test | $\mathcal{V}$ | — | $85.72 \pm 5.32$ |
| FW | SimpleMKL | $\mathcal{V}$ | 168130 | Out of memory |
| | EasyMKL | $\mathcal{V}$ | 168130 | $86.12 \pm 4.54$ |
| FWS | EasyMKLFS | $\mathcal{V}$ | 168130 | $86.91 \pm 5.12$ |

The second step is to combine heterogeneous data (image and non-image features) for prediction. Table 3 shows the results obtained when we combine both image and clinical features in different ways. Combining the MR images with the clinical information by concatenation (i.e. SVM with **I** + **C**) or by using standard MKL or RF approaches produces a model that is similar (in accuracy) to the one generated by using only the MR features. A small improvement of the results is obtained by the feature selection methods (i.e. SVM RFE and SVM t-test). EasyMKL used as feature weighter provides a larger improvement, because it is able to select a single weight for each voxel of the MR image. Finally, by removing the noise from the weights of EasyMKL, the proposed method (EasyMKLFS) is able to provide the best performance.

**Table 3:**
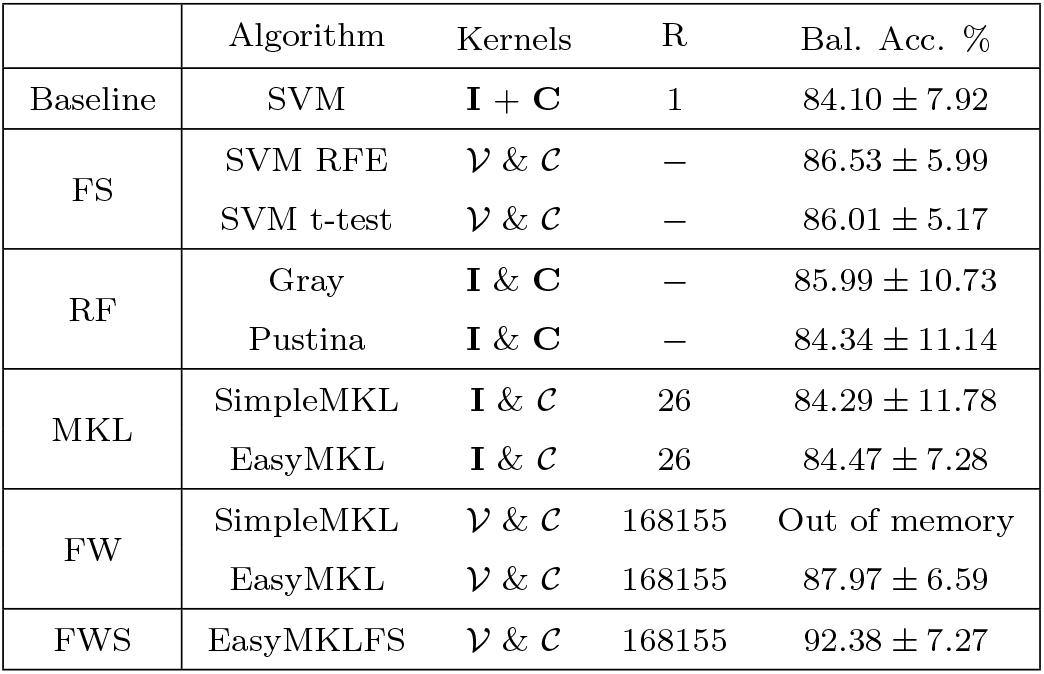
ADNI Dataset: comparisons of 5 repetitions of a nested 10-fold cross-validation balanced accuracy using the clinical information selected by a FDR procedure. The results are divided in 5 families: Baseline, Feature Selection (FS), Random Forests-based family (RF), standard Multiple Kernel Learning (MKL), Feature Weighting by using MKL (FW) and our method in Feature Weighting and Selection (FWS). R corresponds to the number of kernels used.

|  | Algorithm | Kernels | R | Bal. Acc. % |
| --- | --- | --- | --- | --- |
| Baseline | SVM | $\mathbf{I} + \mathbf{C}$ | 1 | $84.10 \pm 7.92$ |
| FS | SVM RFE | $\mathcal{V} \& \mathcal{C}$ | — | $86.53 \pm 5.99$ |
| | SVM t-test | $\mathcal{V} \& \mathcal{C}$ | — | $86.01 \pm 5.17$ |
| RF | Gray | $\mathbf{I} \& \mathbf{C}$ | — | $85.99 \pm 10.73$ |
| | Pustina | $\mathbf{I} \& \mathbf{C}$ | — | $84.34 \pm 11.14$ |
| MKL | SimpleMKL | $\mathbf{I} \& \mathcal{C}$ | 26 | $84.29 \pm 11.78$ |
| | EasyMKL | $\mathbf{I} \& \mathcal{C}$ | 26 | $84.47 \pm 7.28$ |
| FW | SimpleMKL | $\mathcal{V} \& \mathcal{C}$ | 168155 | Out of memory |
| | EasyMKL | $\mathcal{V} \& \mathcal{C}$ | 168155 | $87.97 \pm 6.59$ |
| FWS | EasyMKLFS | $\mathcal{V} \& \mathcal{C}$ | 168155 | $92.38 \pm 7.27$ |

In order to compare the predictions of the proposed EasyMKLFS with respect to the other methods we used the non-parametric Wilcoxon signed-rank test [56]. The results of these tests are presented in Table 4. Since there were 8 comparisons, the Bonferroni corrected p-value is 0.05/8 = 6.25 · 10^−3^. Not surprising the test showed a significance difference between the proposed methods with respect to all compared approaches, and the one with the performance most similar to the EasyMKLFS is the EasyMKL.

**Table 4:** ADNI Dataset: results of the Wilcoxon signed-rank test comparing EasyMKLFS with respect to the others. Smaller p-values mean an higher difference between the models and, in our case, the Bonferoni corrected p-value is 0.05/8 = 6.25 · 10^−3^.

|  | Algorithm | p-value w.r.t. EasyMKLFS |
| --- | --- | --- |
| Baseline | SVM | $2.7 \cdot 10^{-5}$ |
| FS | SVM RFE | $3.2 \cdot 10^{-5}$ |
| | SVM t-test | $5.6 \cdot 10^{-4}$ |
| RF | Gray | $1.9 \cdot 10^{-7}$ |
| | Pustina | $9.1 \cdot 10^{-6}$ |
| MKL | SimpleMKL | $3.8 \cdot 10^{-4}$ |
| | EasyMKL | $3.7 \cdot 10^{-4}$ |
| FW | EasyMKL | $1.7 \cdot 10^{-3}$ |

Figure 2 shows the selection frequency for the FS sparse methods (SVM RFE and SVM t-test) or the average of the weights ***η*** (for EasyMKLFS), respectively, overlaid onto an anatomical brain template, which can be used as a surrogate for consistency. These maps show that all approaches find brain areas previously identified as important for neuroimaging-based diagnosis of Alzheimer (e.g. bilateral hippocampus and amygdala). However, the SVM RFE and SVM t-test also select features across the whole brain potentially related to noise, while the EasyMKLFS selects almost exclusively voxels within the hippocampus and amygdala. In Table 5 we present the top 10 most selected regions by each method (SVM RFE, SVM t-test and EasyMKLFS).

**Figure 2:**
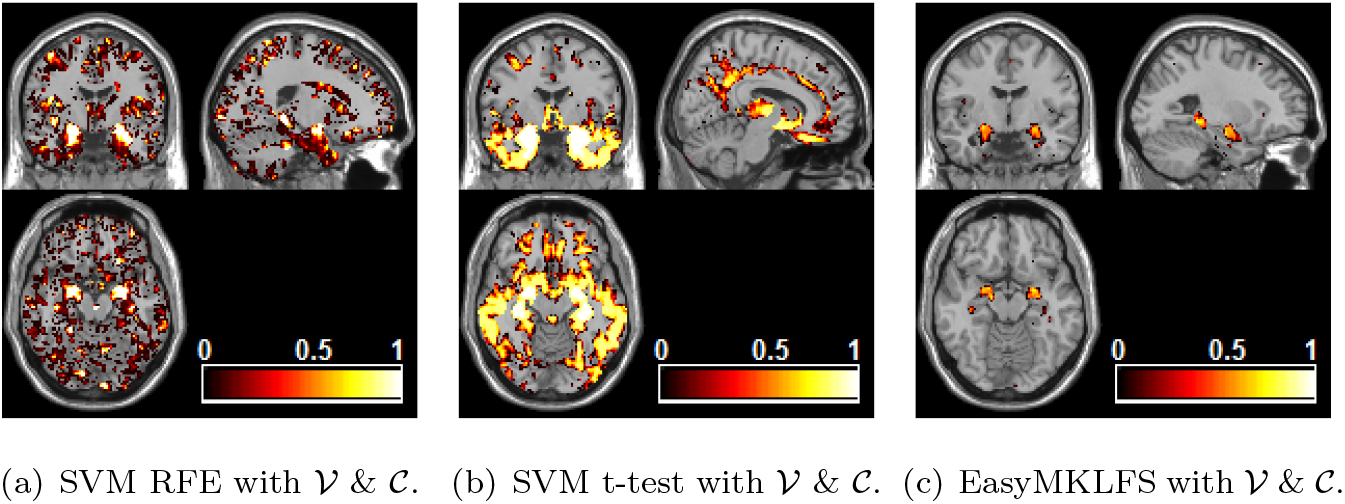
ADNI dataset: comparison of voxels selection frequency (RFE and t-test) and weights (EasyMKLFS), overlayed onto an anatomical template.

**Table 5:** ADNI dataset: the top 10 most selected brain regions for SVM RFE, SVM t-test and EasyMKLFS (with respect to the assigned weight) with the number of selected voxels.

| SVM RFE | voxels | SVM t-test | voxels | EasyMKLFS | voxels |
| --- | --- | --- | --- | --- | --- |
| Amygdala-L | 188 | Amygdala-L | 202 | Amygdala-L | 121 |
| Amygdala-R | 210 | Amygdala-R | 231 | Amygdala-R | 102 |
| Hippocampus-L | 713 | Hippocampus-L | 747 | Hippocampus-L | 255 |
| Hippocampus-R | 659 | ParaHippocampal-L | 798 | Hippocampus-R | 264 |
| ParaHippocampal-L | 738 | Hippocampus-R | 739 | ParaHippocampal-L | 142 |
| ParaHippocampal-R | 725 | ParaHippocampal-R | 877 | ParaHippocampal-R | 88 |
| Temporal-Inf-L | 1844 | Temporal-Inf-L | 2622 | Vermis-4-5 | 30 |
| Vermis-8 | 165 | Fusiform-L | 1734 | Temporal-Inf-L | 118 |
| SupraMarginal-L | 653 | Temporal-Inf-R | 2694 | SupraMarginal-L | 37 |
| Vermis-7 | 110 | Fusiform-R | 1723 | Lingual-L | 32 |

In Figure 3, the weights assigned to the clinical information by EasyMKL are depicted. These weights are generated by using 𝒱 & 𝒞 as family of basic kernels. The top 5 highest weights are assigned to some of the clinical information concerning the MMSE questionnaire, specifically the task related to write a sentence (MMWRITE), put a paper on the floor (MMONFLR), repeat a name of an object (the word “tree” for MMTREE and the word “flag” for MMFLAG) and answer to a simple question about an object (in this case a wrist watch for MMWATCH). See Table B.12 for further information.

**Figure 3:**
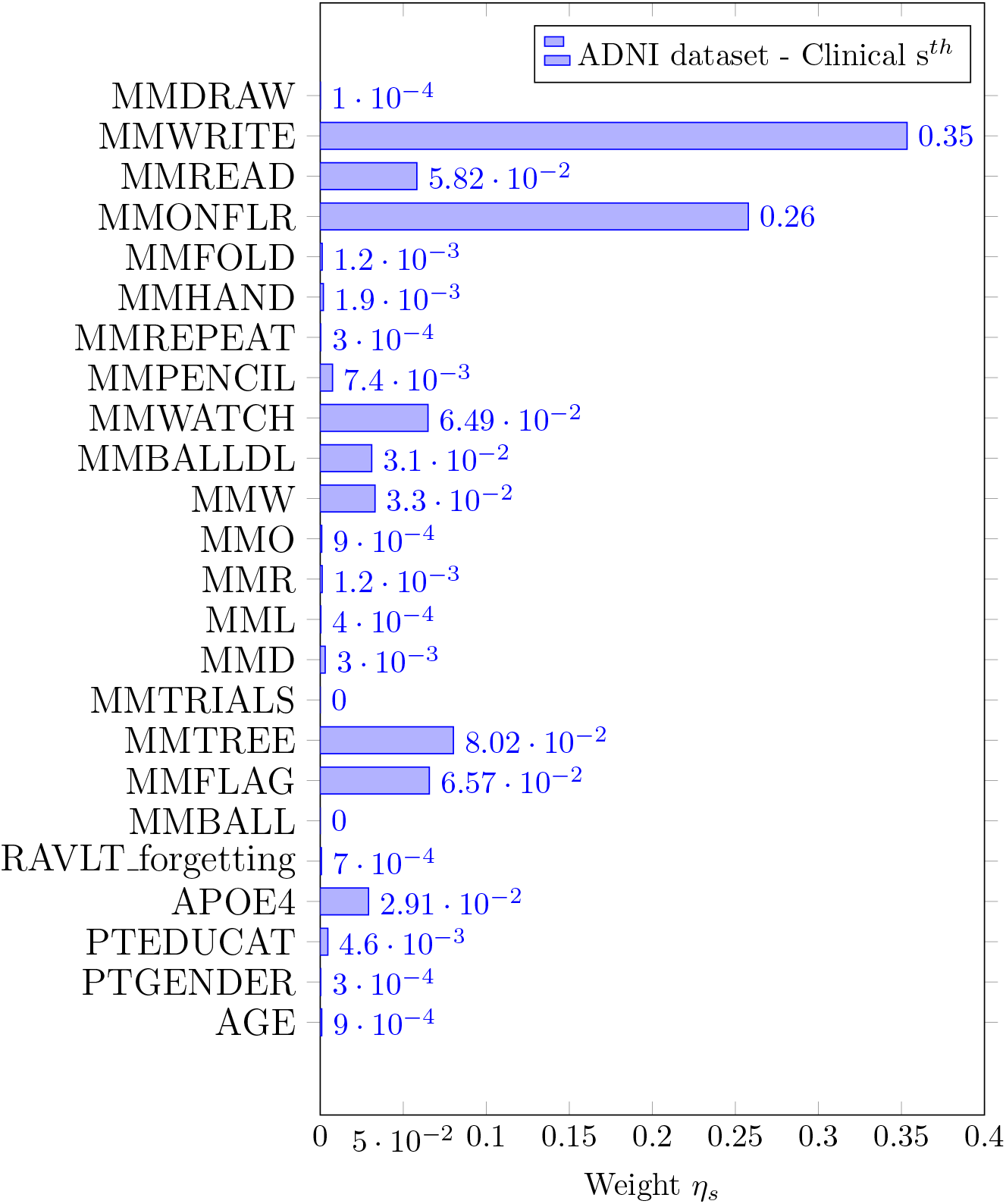
EasyMKL assigned weights for the clinical information selected by a FDR procedure exploiting 𝒱 & 𝒞 as family of basic kernels for the ADNI dataset. The top 5 highest weights are assigned to the clinical data (see Table B.12 for further information): MMWRITE, MMONFLR, MMTREE, MMFLAG and MMWATCH.

Figure 4 depicts the cumulative weight assigned by EasyMKLFS to each source of information (sMRI and clinical information). These weights show that the importance of the sMRI images is larger than the clinical data. Nevertheless, the accuracy results show that the clinical features contributed to the improvement of the final predictive model (changing the performance of our method from 86.91% to 92.38% of balanced accuracy, in this classification task).

**Figure 4:**
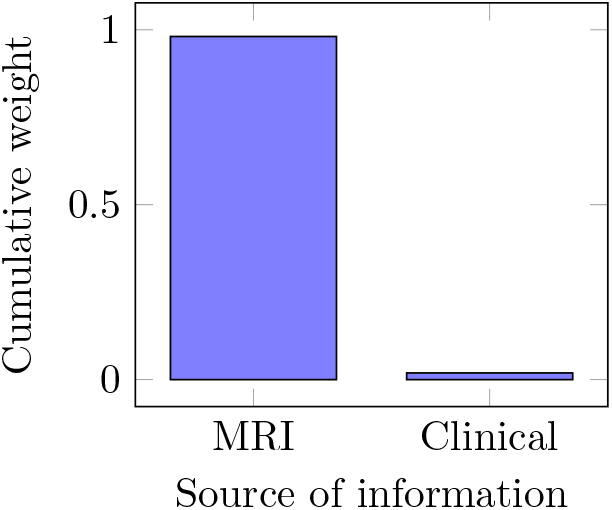
EasyMKLFS assigned weights for the different sources of information: MR images and clinical measurements.

### 4.2. Depression

In this section we present the results obtained by using the Depression dataset. Table 6 shows the results obtained by exploiting each source of information alone, i.e. the clinical data or the combination of the four fMRI derived images of each subject (brain activation to Anxious, Happy, Neutral and Sad faces). These results highlight the challenge of this classification task. In this case, the clinical features bring a good amount of information, which is comparable with the information contained in the fMRI. In fact, the best accuracy of the single source methods is 79.67% for Linear SVM with the clinical data, and 68% with EasyMKL with the fMRIs features. Due to the fact that this dataset includes a very heterogeneous group of patients, the training labels are extremely “noisy” and unreliable. For this reason, the standard feature selection methods (i.e. SVM RFE and SVM t-test) fail to select the relevant voxels. Our method showed a similar performance to EasyMKL (used as a simple feature weighter) but it is able to produce a sparser solution, providing more interpretability when compared with a dense model.

**Table 6:** Depression Dataset: comparisons of 5 repetitions of a nested 10-fold cross-validation balanced accuracy using the clinical information selected by a FDR procedure. The results are divided in 4 families: Baseline, Feature Selection (FS), Feature Weighting by using MKL (FW) and our method in Feature Weighting and Selection (FWS). R corresponds to the number of kernels used.

|  | Algorithm | Kernels | R | Bal. Acc. % |
| --- | --- | --- | --- | --- |
| Baseline | SVM | **C** | 1 | $79.67 \pm 12.29$ |
| | SVM | **I** | 1 | $67.00 \pm 14.87$ |
| FS | SVM RFE | $\mathcal{V}$ | – | $65.33 \pm 12.97$ |
| | SVM t-test | $\mathcal{V}$ | – | $62.19 \pm 10.12$ |
| FW | SimpleMKL | $\mathcal{V}$ | 713816 | Out of memory |
| | EasyMKL | $\mathcal{V}$ | 713816 | $68.00 \pm 13.67$ |
| FWS | EasyMKLFS | $\mathcal{V}$ | 713816 | $67.73 \pm 11.32$ |

Similarly to the previous example, we avoid the comparison of FS or FW methods using only the clinical information, due to the low dimensionality of the problem with respect to the number of the examples.

Table 7 shows the results by combining the fMRI derived features with the clinical information. For this challenging classification task, the FS methods showed similar performance with and without the clinical information. Some improvement is obtained by the RF approaches, however a slightly bigger improvement is provided by the standard MKL methods (with an accuracy of 79.67% for SimpleMKL). The results of the EasyMKL, EasyMKL as FW, and our method (EasyMKLFS), are comparable to standard MKL. However, once again, our method produces a sparse model, which is more interpretable.

**Table 7:** Depression Dataset: comparisons of 5 repetitions of a nested 10-fold cross-validation balanced accuracy using the clinical information selected by a FDR procedure. The results are divided in 5 families: Baseline, Feature Selection (FS), Random Forests-based family (RF), standard Multiple Kernel Learning (MKL), Feature Weighting by using MKL (FW) and our method in Feature Weighting and Selection (FWS). R corresponds to the number of kernels used.

|  | Algorithm | Kernels | R | Bal. Acc. % |
| --- | --- | --- | --- | --- |
| Baseline | SVM | $\mathbf{I} + \mathbf{C}$ | 1 | $67.00 \pm 14.87$ |
| FS | SVM RFE | $\mathcal{V} \ \& \ \mathcal{C}$ | — | $64.99 \pm 13.01$ |
| | SVM t-test | $\mathcal{V} \ \& \ \mathcal{C}$ | — | $62.72 \pm 11.12$ |
| RF | Gray | $\mathbf{I} \ \& \ \mathbf{C}$ | — | $75.34 \pm 16.34$ |
| | Pustina | $\mathbf{I} \ \& \ \mathbf{C}$ | — | $73.88 \pm 15.19$ |
| MKL | SimpleMKL | $\mathbf{I} \ \& \ \mathcal{C}$ | 45 | $79.67 \pm 13.11$ |
| | EasyMKL | $\mathbf{I} \ \& \ \mathcal{C}$ | 45 | $79.61 \pm 14.12$ |
| FW | SimpleMKL | $\mathcal{V} \ \& \ \mathcal{C}$ | 713860 | Out of memory |
| | EasyMKL | $\mathcal{V} \ \& \ \mathcal{C}$ | 713860 | $80.02 \pm 11.32$ |
| FWS | EasyMKLFS | $\mathcal{V} \ \& \ \mathcal{C}$ | 713860 | $80.01 \pm 10.11$ |

As for the ADNI dataset, we compared the different methods with respect the proposed EasyMKLFS concerning the predictions performing the non-parametric Wilcoxon signed-rank test [56]. The results of the p-values obtained from of these tests are presented in Table 8. Similarly to the previous dataset the Bonferroni corrected p-value is 0.05/8 = 6.25 · 10^−3^. The differences are significant for all the methods but EasyMKL. EasyMKL is a fundamental part of the proposed algorithm. EasyMKLFS combines the properties of EasyMKL with feature selection. The uncertainty of the labels and the amount of noise in the Depression dataset probably makes the feature selection step not as beneficial as in the previous example.

**Table 8:** Depression Dataset: results of the Wilcoxon signed-rank test comparing EasyMKLFS with respect to the others. Smaller p-values mean an higher difference between the models and, in our case, the Bonferoni corrected p-value is 0.05/8 = 6.25 · 10^−3^.

|  | Algorithm | p-value w.r.t. EasyMKLFS |
| --- | --- | --- |
| Baseline | SVM | $8.6 \cdot 10^{-5}$ |
| FS | SVM RFE | $3.8 \cdot 10^{-4}$ |
| | SVM t-test | $1.2 \cdot 10^{-4}$ |
| RF | Gray | $4.3 \cdot 10^{-5}$ |
| | Pustina | $7.8 \cdot 10^{-4}$ |
| MKL | SimpleMKL | $1.8 \cdot 10^{-4}$ |
| | EasyMKL | $4.6 \cdot 10^{-4}$ |
| FW | EasyMKL | $9.6 \cdot 10^{-3}$ |

Figure 7 shows the selection frequency of the sparse FS methods (SVM RFE and SVM t-test) or the average of the weights ***η*** (for EasyMKLFS) overlaid onto an anatomical brain template, which can be used as a surrogate of consistency. For each method, we present the selection frequency or the average of the weights for the four fMRI derived images (i.e. brain activation to Anxious, Happy, Neutral and Sad faces). In Tables 9 and 10, we present the top 10 brain regions selected for each method (SVM RFE, SVM t-test and EasyMKLFS), and for each fMRI derived image. The vast majority of these regions has been previously described in the depression literature. Especially frontal and temporal areas, as well as subcortical regions, such as: the hippocampus, the amygdala, and parts of the reward system (e.g. the pallidum and the caudate). These regions have been previously identified using both multivariate pattern recognition approaches, and classic group statistical analyses [23, 60–62].

**Table 9:** Depression dataset: the top 10 most selected brain regions for SVM RFE, SVM t-test and EasyMKLFS (with respect to the assigned weight) with the number of selected voxels.

| Anxious |  |  |  |  |  |
| --- | --- | --- | --- | --- | --- |
| SVM RFE | voxels | SVM t-test | voxels | EasyMKLFS | voxels |
| Calcarine-L | 783 | Pallidum-L | 147 | Calcarine-L | 266 |
| Occipital-Sup-R | 364 | Putamen-L | 514 | Temporal-Sup-L | 237 |
| Frontal-Sup-Medial-R | 692 | SupraMarginal-R | 874 | Occipital-Sup-R | 134 |
| Calcarine-R | 532 | Occipital-Sup-R | 517 | Paracentral-Lobule-R | 82 |
| Temporal-Sup-L | 655 | Postcentral-R | 1617 | Frontal-Mid-L | 394 |
| Parietal-Sup-L | 534 | Frontal-Mid-L | 1793 | Frontal-Sup-R | 319 |
| Frontal-Mid-L | 1104 | Paracentral-Lobule-R | 296 | Frontal-Sup-Medial-R | 157 |
| Paracentral-Lobule-R | 224 | Calcarine-R | 662 | Frontal-Inf-Tri-L | 184 |
| Temporal-Sup-R | 869 | Frontal-Sup-R | 1395 | Frontal-Inf-Oper-L | 69 |
| Cingulum-Mid-L | 629 | Cuneus-R | 481 | Temporal-Mid-R | 318 |
| Happy |  |  |  |  |  |
| SVM RFE | voxels | SVM t-test | voxels | EasyMKLFS | voxels |
| SupraMarginal-L | 358 | Cingulum-Ant-R | 630 | Temporal-Sup-L | 366 |
| Temporal-Sup-L | 666 | Cuneus-R | 587 | SupraMarginal-L | 204 |
| Calcarine-L | 805 | Temporal-Sup-L | 677 | Paracentral-Lobule-R | 73 |
| Precentral-R | 1063 | Hippocampus-L | 311 | Calcarine-L | 222 |
| Insula-R | 335 | Putamen-R | 388 | Precentral-R | 268 |
| Putamen-R | 216 | Hippocampus-R | 449 | Insula-R | 128 |
| Temporal-Mid-R | 1339 | Calcarine-R | 715 | Putamen-R | 64 |
| Caudate-R | 295 | Caudate-L | 548 | Frontal-Mid-L | 365 |
| Caudate-L | 331 | Thalamus-L | 500 | Caudate-L | 87 |
| Calcarine-R | 474 | Cuneus-L | 652 | Frontal-Sup-R | 311 |

**Table 10:** Depression dataset: the top 10 most selected Atlas Regions of the brain for SVM RFE, SVM t-test and EasyMKLFS (with respect to the assigned weight) with the number of selected voxels.

| Neutral |  |  |  |  |  |
| --- | --- | --- | --- | --- | --- |
| SVM RFE | voxels | SVM t-test | voxels | EasyMKLFS | voxels |
| Temporal-Sup-L | 775 | Hippocampus-L | 447 | Temporal-Sup-L | 398 |
| Amygdala-R | 115 | Thalamus-L | 624 | SupraMarginal-L | 175 |
| Temporal-Mid-R | 1444 | Hippocampus-R | 547 | Pallidum-R | 44 |
| SupraMarginal-L | 388 | Amygdala-R | 131 | Amygdala-L | 26 |
| Amygdala-L | 79 | Temporal-Sup-L | 877 | Thalamus-L | 129 |
| Thalamus-L | 461 | Putamen-R | 488 | Temporal-Mid-R | 470 |
| Pallidum-R | 97 | Putamen-L | 426 | Hippocampus-L | 82 |
| Hippocampus-R | 329 | Temporal-Mid-R | 1618 | Hippocampus-R | 86 |
| Caudate-R | 299 | Caudate-R | 441 | Putamen-R | 75 |
| Hippocampus-L | 238 | ParaHippocampal-L | 359 | Precentral-R | 260 |
| Sad |  |  |  |  |  |
| SVM RFE | voxels | SVM t-test | voxels | EasyMKLFS | voxels |
| Parietal-Sup-L | 717 | Amygdala-R | 117 | Temporal-Sup-L | 342 |
| Temporal-Sup-L | 760 | Postcentral-R | 1462 | SupraMarginal-L | 159 |
| SupraMarginal-L | 383 | Cingulum-Ant-R | 554 | Precentral-R | 310 |
| Precentral-R | 986 | Temporal-Sup-L | 783 | Parietal-Sup-L | 199 |
| Caudate-L | 213 | Caudate-L | 398 | Caudate-L | 87 |
| Insula-L | 506 | Parietal-Sup-L | 934 | ParaHippocampal-L | 69 |
| Thalamus-L | 313 | Hippocampus-L | 342 | ParaHippocampal-R | 68 |
| Temporal-Pole-Sup-L | 269 | Occipital-Sup-R | 598 | Insula-L | 122 |
| Postcentral-R | 768 | Frontal-Mid-L | 1625 | Frontal-Inf-Tri-L | 150 |
| Occipital-Mid-R | 556 | Putamen-R | 303 | Frontal-Mid-L | 256 |

Figure 5 depicts the weights assigned by EasyMKL for the clinical information. The family 𝒱 & 𝒞 has been used for the basic kernels. For this dataset, the top 5 highest weights are assigned to the following clinical information: the Negative Affect Schedule (PANAS_neg), the mean valence ratings for male neutral and sad faces (from KDEF, i.e. KDEF_val_neu_m and KDEF_val_sad_m), the mean arousal rating for male happy faces (from KDEF, i.e. KDEF_aro_hap_m) and an extracted feature from the State-Trait anger expression inventory test (STAXI_TAT). See Table B.13 for further information.

**Figure 5:**
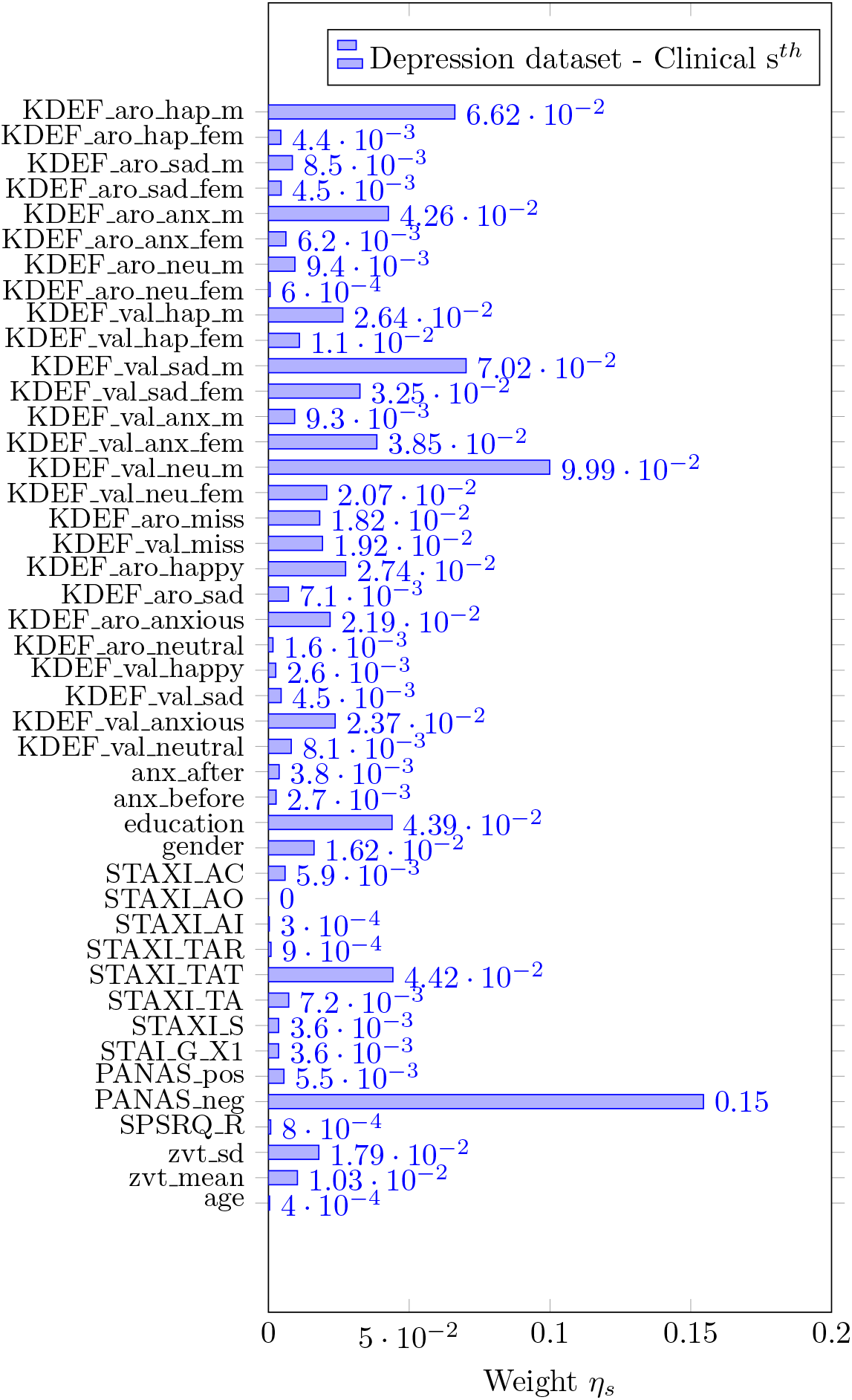
EasyMKL assigned weights for the clinical information selected by a FDR procedure exploiting 𝒱 & 𝒞 as family of basic kernels for the Depression dataset. The top 5 highest weights are assigned to the clinical data (see Table B.13 for further information): PANAS_neg, KDEF_val_neu_m, KDEF_val_sad_m, KDEF_aro_hap_m and STAXI_TAT.

Figure 6 shows the sums of the weights that are assigned for each information source (4 fMRI derived images plus the clinical information) by our method.

**Figure 6:**
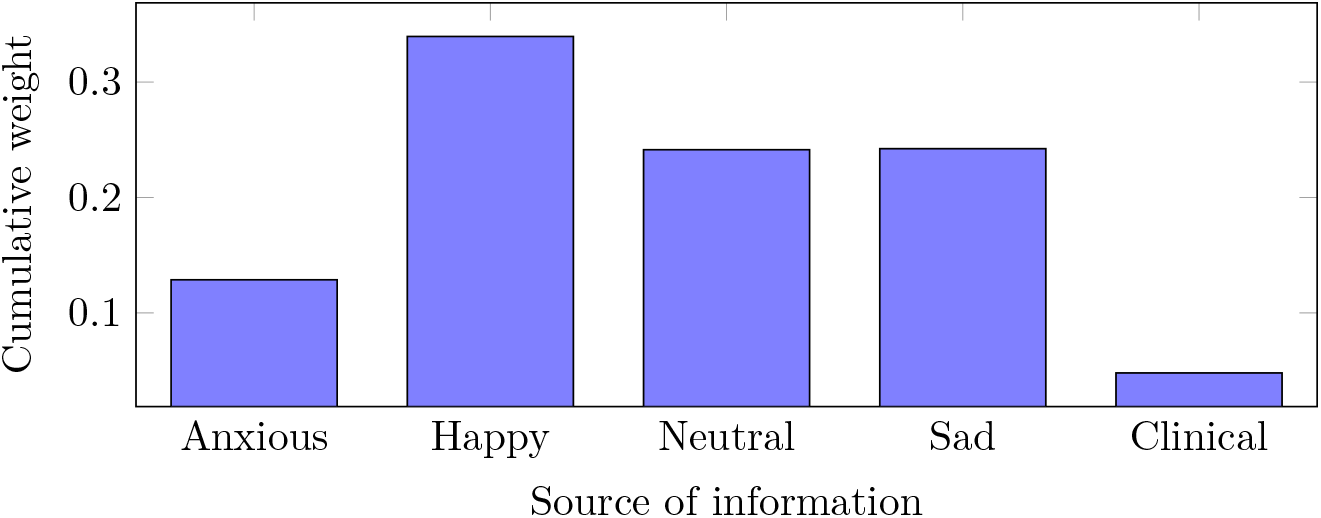
EasyMKLFS assigned weights for the different sources of information of the Depression dataset: Anxious image, Happy image, Neutral image, Sad image and clinical measurements.

**Figure 7:**
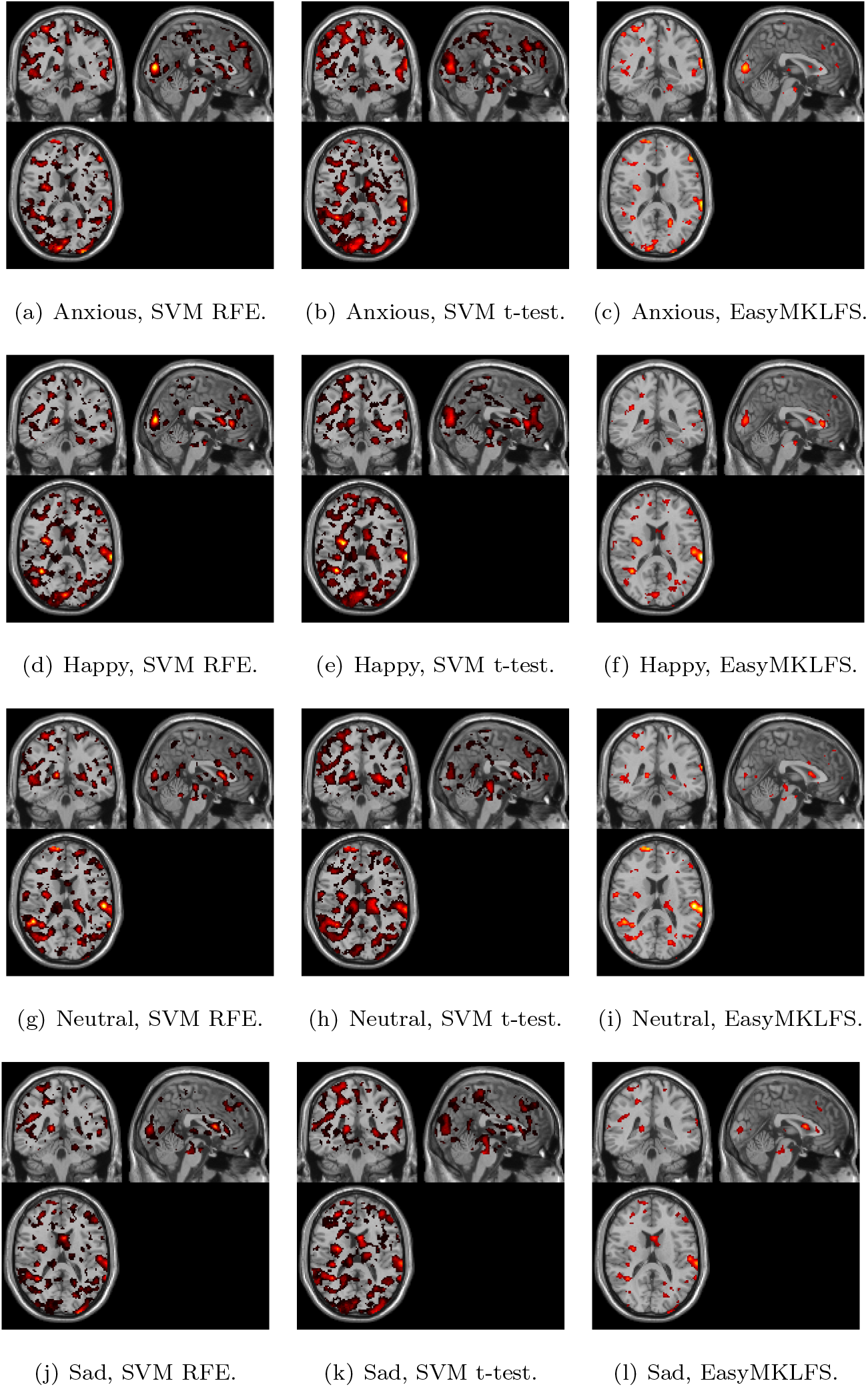
Depression dataset: comparison of voxels selection frequency (RFE and t-test) and weights (EasyMKLFS) by using 𝒱 & 𝒞, overlayed onto an anatomical template.

## 5. Discussion

The main goal of this paper is to present an effective methodology to combine and select features from different sources of information (sMRI/fMRI, clinical and demographic information) in order to classify patients with mental health disorders versus healthy controls. The proposed method (EasyMKLFS) obtained better or similar accuracy than several compared machine learning approaches with higher levels of sparsity, therefore consistently improving interpretability.

More specifically, by using the ADNI dataset, we were able to obtain a significant improvement in the classification accuracy, potentially due to absence of strong source of noise in the data and presence of predictive information in the considered sources of information. On the other hand, in the Depression dataset, we obtained a comparable accuracy to the MKL gold standard methods. The lack of a significant improvement in classification accuracy for the depression dataset might be explained by the noise in the fMRI data and higher label uncertainty for this task (i.e. high heterogeneity in the depressed group). More important, in both the cases, the EasyMKLFS provides the sparser solution. This particular result improves the interpretability of our models, identifying which features are driving the predictions.

In the context of machine learning, interpretability of a model often refers to its ability to identify a subset of informative features. In contrast, in neuroscience and clinical neuroscience, researchers often wants to understand why a specific feature contribute or is informative to a predictive model. Unfortunately, answering the question of why a feature is informative to a predictive model is not straightforward and has been topic of a number of studies in the field of neuroimaging (e.g. [63–66]). These studies have shown that a features can be included in a model due to different reasons (e.g. a feature might be informative because it has consistently high/low value for one class with respect to the other class or because it helps canceling correlated noise). In the present work we use the machine learning definition of model interpretability or informativeness. The identified features were compared with previous literature in terms of how they overlap with regions previously described as important for discriminating dementia and depression from healthy subjects.

It is important to note what makes our method different from the standard approaches to combine heterogeneous information for neuroimaging based diagnosis. EasyMKLFS works in a framework where the initial information is fragmented in small and low informative pieces, and without exploiting some *a priori* knowledge from an expert. Due to the particular ability of EasyMKL to combine huge amounts of different kernels (i.e. one per feature), we are able to weight all of them. This first difference with respect to the state-of-art MKL applications is crucial, in fact, other MKL methods often combine only a small set of different sources manually selected. Our method is able to work without this bias and obtain better or similar performance as previous methods. Finally, the last step of EasyMKLFS is able to find a very sparse model unifying in synergy the characteristics of feature weighting (i.e. MKL with a large amount of basic kernels) and feature selection.

When compared to the RF-based approaches, our method obtains better accuracy and, as in the MKL case, the main difference is the computational complexity of these methods. In fact, the two RF-based methodologies (i.e. Pustina and Gray) have an increase in computational time to perform the training that is orders of magnitude higher when the number of different sources of information increase. Moreover, these approaches are a mixture of heuristics and algorithms, not easily comparable to the other well-theoretically-grounded machine learning methods used in the paper.

In our experiments, we reported the average accuracy of each method together with its standard deviation. This procedure is broadly used comparing machine learning methods. For the sake of completeness, we have compared the performance of the proposed algorithm, EasyMKLFS, with each of the other methods using the Wilcoxon signed-rank test [56]. Results from these comparisons show that the EasyMKLFS was significantly better than all other methods for the ADNI dataset and significantly better than all but the EasyMKL for the depression dataset. The lack of improvement with respect to the EasyMKL for the depression dataset suggests that for heterogeneous datasets with high label uncertainty (i.e. datasets that contain subgroups of subjects with different characteristics) the feature selection step might not be advantageous. Unfortunately, label uncertainty is a common issue in psychiatry disorders. Current diagnostic categories in psychiatric are only based on symptoms and behaviours due to the lack of biomarkers in psychiatry [67]. There is a lot of evidence that the boundary of these categories do not alight with neuroscience, genetics and have also not been predictive of treatment response [68]. Another evidence of the impact of class heterogeneity on the performance of neuroimaging based classifiers can be found in [69] where the author shows a negative correlation between reported accuracy and sample size for many diagnostic applications. Bigger samples are likely to be more heterogeneous than small ones. In summary, taken together, these results demonstrate the effectiveness of our methodology in two different classification tasks, obtaining similar or higher accuracy than the compared methods with higher interpretability.

The EasyMKLFS was able to identify, for both datasets, sMRI/fMRI and clinical/demographic features that overlap with features previously identified as relevant for discriminating demented and depressed patients from healthy controls. More specifically, for the ADNI dataset, the top most selected brain regions according to the AAL atlas were bilateral amygdala, hippocampus and parahippocampus. The top most selected clinical information were items of the Mini-Mental State Examination (MMSE).The MMSE is a 30-point questionnaire that is used extensively in clinical and research settings to measure cognitive impairment [45]. The depression dataset consisted of four brain images, representing fMRI patterns of brain activation to different emotional faces (Anxious, Happy, Neutral and Sad), in addition to the clinical information. The top most selected brain regions across the different emotions included frontal and temporal areas, as well as subcortical regions, such as: the hippocampus, the amygdala, and parts of the reward system (e.g. the pallidum and the caudate). All these regions have been has been previously described in the depression literature [23, 60–62]. The top most selected clinical information for the depression dataset was the Negative Affect Schedule (PANAS neg). The Positive and Negative Affect Schedule (PANAS) is a self-report questionnaire that measures both positive and negative affect [70]. Previous studies have shown that individuals with higher Negative Affect (NA) trait (neuroticism) show heightened emotional reactivity [71] and experience more negative emotions [72]. Higher NA trait has been also associated with poor prognosis [72] and predictive of onset of major depression [73]. Furthermore, a recent study showed that it is possible to decode individuals NA trait from patterns of brain activation to threat stimuli in a sample of healthy subject [74]. Our results, corroborate with these previous studies and support the evidence that Negative Affect trait might have important clinical implications for depression.

From a clinical perspective, the proposed approach addresses the two fundamental challenges arising from the unique, multivariate and multi-modal nature of mental disorders (for an in-depth discussion of both conceptual challenges, see [75]). On the one hand, mental disorders are characterized by numerous, possibly interacting biological, intrapsychic, interpersonal and socio-cultural factors [76, 77]. Thus, a clinically useful patient representation must, in many cases, include data from multiple sources of observation, possibly spanning the range from molecules to social interaction. Even within the field of neuroimaging, we see a plethora of modalities used in daily research; including e.g. task-related and resting-state fMRI, structural MRI data and Diffusion Tensor Imaging (DTI) approaches. All these modalities might contain non-redundant, possibly interacting sources of information with regard to the clinical question. In fact, it is this peculiarity – distinguishing psychiatry from most other areas of medicine – which has hampered research in general and translational efforts for decades. Overwhelming evidence shows that no single measurement – be it a voxel, a gene or a psychometric test – explains substantial variance with regards to any practically relevant aspect of a psychiatric disorder (compare e.g. [78]). In addition, many if not most variables are irrelevant for the particular question addressed. It is this profoundly multivariate nature of mental disorders that necessitates dimensionality reduction or feature-selection approaches when using whole-brain neuroimaging data. The fact that EasyMKLFS now addresses, both, the issue of feature selection and multi-modal data integration in a single, mathematically principled framework constitutes a major step forward. From a health economic point of view, approaches such as this one are especially noteworthy, as they have the potential not only to identify the best-performance, but also the most efficient model. By using EasyMKLFS, it is possible to directly test which sources of information are non-redundant with regards to the model’s performance.

From the perspective of biomarker research, it is particularly important that EasyMKLFS provides a means to investigate and visualize the predictive model. Using MKL weights in combination with feature selection provides information regarding feature importance for single features, as well as for data sources, while guaranteeing sparsity. Our results show that, compared for example to a classic t-test, the visualization appears much less noisy and focused, dramatically increasing interpretability. Accordingly, we were able to identify many of the key-regions known to be involved in the mental diseases while maintaining a rather focused list of areas.

Despite our encouraging results, the method does present some limitations. Firstly, our method was not able to show an improvement in performance when the classification task is very noisy (i.e. for unreliable patients’ labels), as in the Depression dataset. Another weak point of the presented methodology is that, in this paper, we studied only the simplest way to combine the information, by generating exclusively linear kernels. From this point of view, this is a limitation of our framework with respect to the strength of the kernels methods.

Considering these limitations, there are two possible future directions. Firstly, the improvement of EasyMKL by using a different regularizer that is more stable with respect to the heterogeneity in the patient group. The idea is to split the regularization in two different parts: the first part for the positive examples, and the second part for the negative examples. In this way, we might be able to handle classification with heterogeneous classes better (e.g. the Depression dataset). A second way to evolve our framework is to fragment and to randomly generate the source of information, improving the accuracy by injecting non-linearity. In this sense, a good way to proceed is by randomly generating small subsets of information from the raw data, then projecting them onto a non-linear feature space before the weighting and selection phase. In this way, we might be able to increase the expressiveness of our features and, consequently, the complexity of the generated model. On the other hand, we have to be able to bound these new degrees of freedom, in order to avoid overfitting.

In terms of future applications, the proposed EasyMKLFS approach has the ability to be applied to other clinical relevant classification tasks such as distinguishing diseases groups and predicting diseases progression (see for example [79–81]). As shown in our results, the performance of the EasyMKLFS approach on these applications will be bounded by the reliability of the labels and informativeness of the considered sources of information. Moreover, our approach might be also particular beneficial for ‘big-data’ applications focusing on personalized medicine, where the goal is to predict future outcomes and/or treatment response by combining larger sources of patient information.

## Acknowledgements

Janaina Mourao-Miranda was funded by the Wellcome Trust under grant number WT102845/Z/13/Z. João M. Monteiro was funded by a PhD scholarship awarded by Fundacao para a Ciencia e a Tecnologia (SFRH/BD/88345/).

## Appendix A. A brief introduction to EasyMKL

As introduced in Section 2.2, EasyMKL [22] is a very efficient MKL algorithm with the clear advantage of having high scalability with respect to the number of kernels to be combined. In fact, its computational complexity is constant in memory and linear in time.

Technically, EasyMKL finds the coefficients ***η*** that maximize the margin on the training set. The margin is computed as the distance between the smaller convex envelopes (i.e. convex hulls) of positive and negative examples in the feature space, as shown in Figure A.8.

**Figure A.8:**
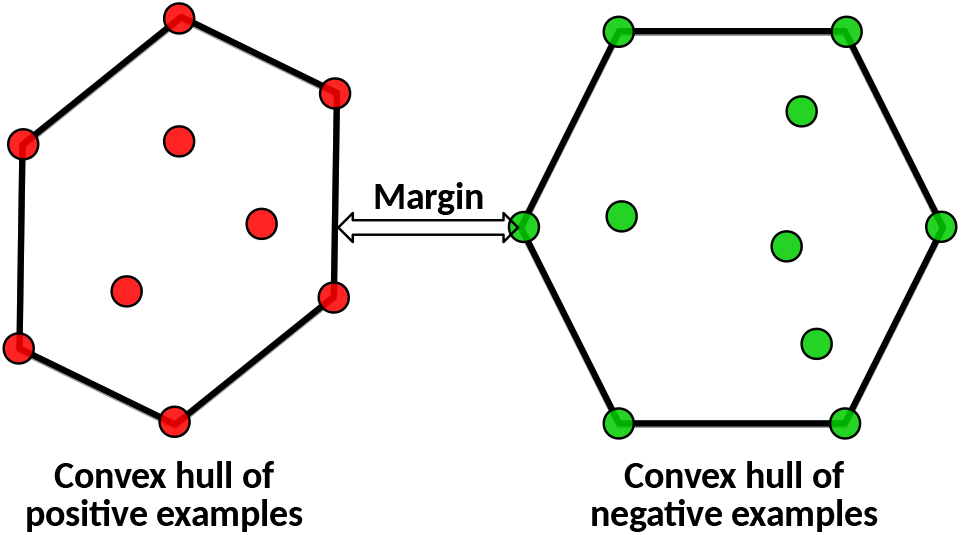
The margin is the distance between the convex hull of the positive examples (in red) and the convex hull of the negative examples (in green). EasyMKL is able to find a combination of kernels that maximizes this distance.

In particular, EasyMKL tries to optimize the following general problem:

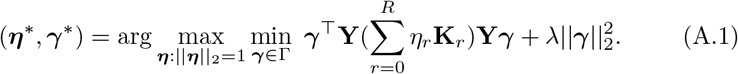

where **Y** is a diagonal matrix with training labels on the diagonal, and λ is a regularization hyper-parameter. The domain Γ represents two probability distributions over the set of positive and negative examples of the training set, that is 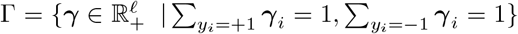. Note that any element ***γ*** ∈ Γ corresponds to a pair of points, the first contained in the convex hull of positive training examples and the second in the convex hull of negative training examples. At the solution, the first term of the objective function represents the obtained (squared) margin, that is the (squared) distance between a point in the convex hull of positive examples and a point in the convex hull of negative examples, in the considered feature space.

Eq. A.1 can be seen as a minimax problem that can be reduced to a simple quadratic problem with a technical derivation described in [22]. The solution of the quadratic problem is an approximation of the optimal ***γ***\* for the original formulation and due to the particular structure of this approximated problem, it is sufficient to provide the average kernel of all the trace-normalized basic kernels, i.e.

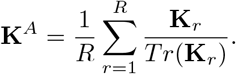

For this reason, we can avoid to store in memory all the single basic kernels obtaining a very scalable MKL algorithm (with respect to the number of kernels).

Finally, from ***γ***\*, it is easy to obtain the optimal weights for the single basic kernels K_r_ by using the following formula

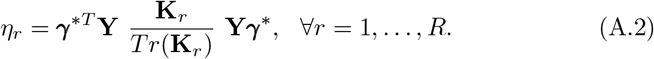

## Appendix B. A further analysis of ADNI and Depression datasets

In Table B.11, the required memory of the different MKL methods is presented. As already noted, SimpleMKL requires a huge amount of memory to handle large family of basic kernels. For example, generating one linear kernel for each voxel, we have to provide more than 50 Gb of memory to store all the required information. EasyMKL and our EasyMKLFS use a fixed amount of memory independently with respect to the number of kernels, due to the particular definition of the optimization problem (see Sections 2.2 and 2.4).

Finally, the list of the extracted clinical information from the ADNI and Depression datasets are summarized in Table B.12 and Table B.13 respectively.

**Table B.11:** ADNI dataset: required memory for different methods to handle different families of basic kernels.

|  | Algorithm | R | Memory | Memory (real) |
| --- | --- | --- | --- | --- |
| Baseline | Linear SVM | 1 | $\mathcal{O}(\ell^2)$ | 293 Kb |
| FS | SVM RFE | – | $\mathcal{O}(\ell^2)$ | 293 Kb |
| | SVM t-test | – | $\mathcal{O}(\ell^2)$ | 293 Kb |
| MKL | SimpleMKL | 26 | $\mathcal{O}(R\ell^2)$ | $\sim 10$ Mb |
| | EasyMKL | 26 | $\mathcal{O}(\ell^2)$ | 293 Kb |
| FW | SimpleMKL | 168155 | $\mathcal{O}(R\ell^2)$ | $\sim 50$ Gb |
| | EasyMKL | 168155 | $\mathcal{O}(\ell^2)$ | 293 Kb |
| FWS | EasyMKLFS | 168155 | $\mathcal{O}(\ell^2)$ | 293 Kb |

**Table B.12:**
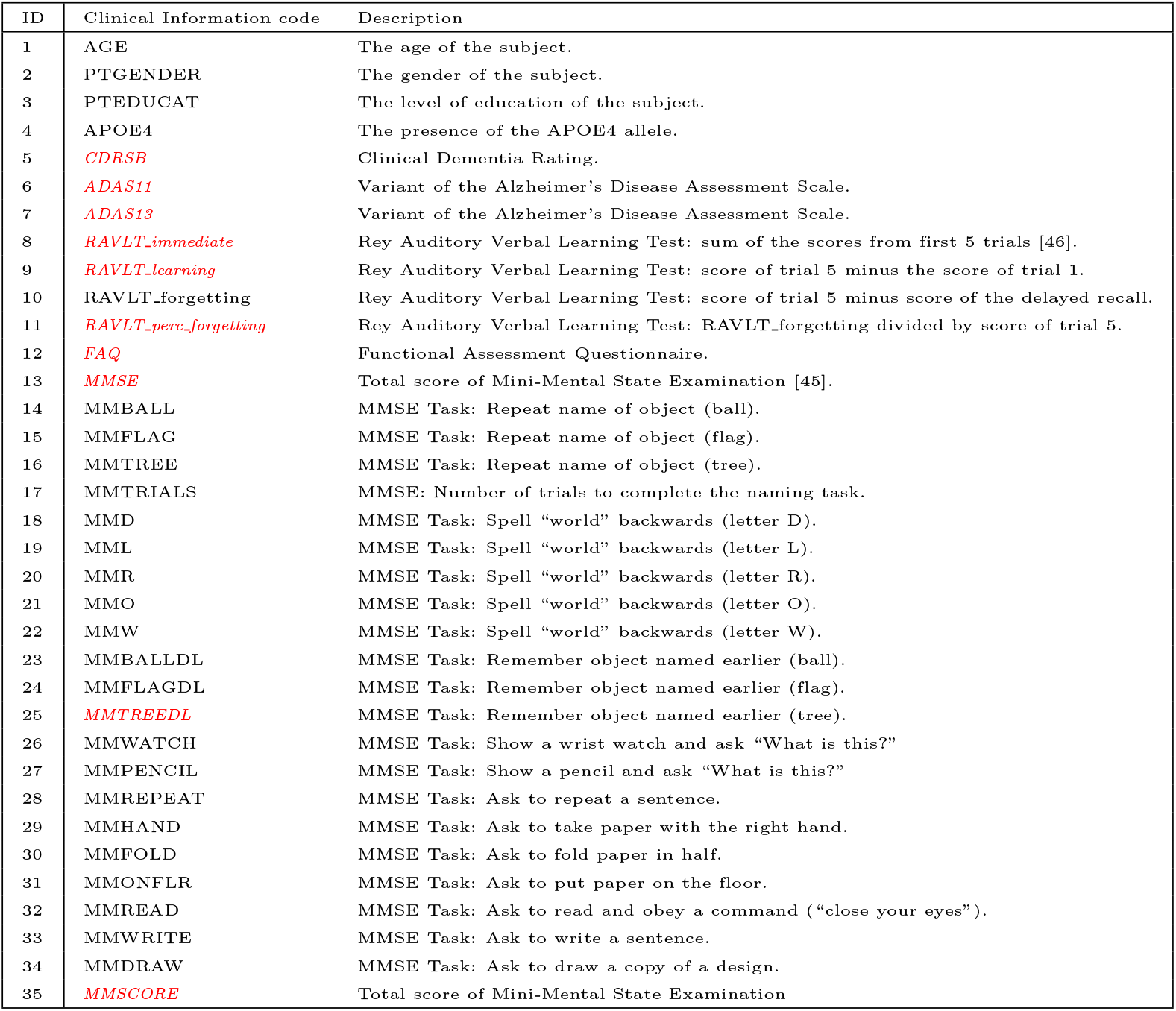
ADNI clinical information. In *italic red*, the clinical information removed by the FDR procedure. All the clinical information starting with “MM” are part of a quite widely used exam that is performed on patients with dementia [45].

**Table B.13:** Depression clinical information. In *italic red*, the clinical information removed by the FDR procedure.

| ID | Clinical Information code | Description |
| --- | --- | --- |
| 1 | age | The age of the patient |
| 2 | zvt_mean | Average of all the tests of processing speed (approx. IQ) [50] |
| 3 | zvt_sd | Standard deviation of all the tests of processing speed |
| 4 | *BDI-II* | Self-report questionnaire of depression severity [52] |
| 5 | *MADRS* | Montgomery-Asberg depression rating scale [51] |
| 6 | SPSRQ_R | Reward score of "Sensitivity to Punishment/Reward Questionnaire" [49] |
| 7 | *SPSRQ_P* | Punishment score of "Sensitivity to Punishment/Reward Questionnaire" |
| 8 | PANAS_neg | Negative Affect Schedule [53] |
| 9 | PANAS_pos | Positive Affect Schedule |
| 10 | STAI_G_X1 | } State-Trait anxiety inventory [54] |
| 11 | *STAI_G_X2* |  |
| 12 | STAXI_S | } State-Trait anger expression inventory [82] |
| 13 | STAXI_TA |  |
| 14 | STAXI_TAT |  |
| 15 | STAXI_TAR |  |
| 16 | STAXI_AI |  |
| 17 | STAXI_AO |  |
| 18 | STAXI_AC |  |
| 19 | gender | The gender of the patient |
| 20 | education | The education level of the patient |
| 21 | anx_before | Visual analog scale of subjective anxiety |
| 22 | anx_after | Anxiety after the scanning |
| 23 | KDEF_val_neutral | Mean Valence ratings for neutral faces from the KDEF [48] collection |
| 24 | KDEF_val_anxious | Mean Valence ratings for Anxious faces from the KDEF collection |
| 25 | KDEF_val_sad | Mean Valence ratings for Sad faces from the KDEF collection |
| 26 | KDEF_val_happy | Mean Valence ratings for Happy faces from the KDEF collection |
| 27 | KDEF_aro_neutral | Mean Arousal ratings for Neutral faces from the KDEF collection |
| 28 | KDEF_aro_anxious | Mean Arousal ratings for Anxious faces from the KDEF collection |
| 29 | KDEF_aro_sad | Mean Arousal ratings for Sad faces from the KDEF collection |
| 30 | KDEF_aro_happy | Mean Arousal ratings for Happy faces from the KDEF collection |
| 31 | KDEF_val_miss | Mean Valence rating missing from the KDEF collection |
| 32 | KDEF_aro_miss | Mean Arousal rating missing from the KDEF collection |
| 33 | KDEF_val_neu_fem | Mean Valence ratings for female Neutral faces from the KDEF collection |
| 34 | KDEF_val_neu_m | Mean Valence ratings for male Neutral faces from the KDEF collection |
| 35 | KDEF_val_anx_fem | Mean Valence ratings for female Anxious faces from the KDEF collection |
| 36 | KDEF_val_anx_m | Mean Valence ratings for male Anxious faces from the KDEF collection |
| 37 | KDEF_val_sad_fem | Mean Valence ratings for female Sad faces from the KDEF collection |
| 38 | KDEF_val_sad_m | Mean Valence ratings for male Sad faces from the KDEF collection |
| 39 | KDEF_val_hap_fem | Mean Valence ratings for female Happy faces from the KDEF collection |
| 40 | KDEF_val_hap_m | Mean Valence ratings for male Happy faces from the KDEF collection |
| 41 | KDEF_aro_neu_fem | Mean Arousal ratings for female Neutral faces from the KDEF collection |
| 42 | KDEF_aro_neu_m | Mean Arousal ratings for male Neutral faces from the KDEF collection |
| 43 | KDEF_aro_anx_fem | Mean Arousal ratings for female Anxious faces from the KDEF collection |
| 44 | KDEF_aro_anx_m | Mean Arousal ratings for male Anxious faces from the KDEF collection |
| 45 | KDEF_aro_sad_fem | Mean Arousal ratings for female Sad faces from the KDEF collection |
| 46 | KDEF_aro_sad_m | Mean Arousal ratings for male Sad faces from the KDEF collection |
| 47 | KDEF_aro_hap_fem | Mean Arousal ratings for female Happy faces from the KDEF collection |
| 48 | KDEF_aro_hap_m | Mean Arousal ratings for male happy Faces from the KDEF collection |

### Supplementary Material

In this section we present the results of the same experiments described in the main paper with the difference that the algorithms can use all the clinical information without any restriction. In the following the results for both the datasets, i.e. ADNI and Depression.

#### Appendix B.1. ADNI

The accuracy results for the ADNI dataset are presented in Table B.14.

**Table B.14:** ADNI Dataset: comparisons of 5 repetitions of a nested 10-fold cross-validation balanced accuracy using all the clinical information. The results are divided in 5 families: Baseline, Feature Selection (FS), standard Multiple Kernel Learning (MKL), Feature Weighting by using MKL (FW) and our method in Feature Weighting and Selection (FWS). R corresponds to the number of kernels used.

|  | Algorithm | Kernels | R | Bal. Acc. % |
| --- | --- | --- | --- | --- |
| Baseline | Linear SVM | $\mathbf{C}$ | 1 | $68.73 \pm 9.68$ |
| | SVM | $\mathbf{I} + \mathbf{C}$ | 1 | $84.80 \pm 6.87$ |
| FS | SVM RFE | $\mathcal{V} \ \& \ \mathcal{C}$ | — | $86.93 \pm 4.76$ |
| | SVM t-test | $\mathcal{V} \ \& \ \mathcal{C}$ | — | $86.47 \pm 6.92$ |
| MKL | SimpleMKL | $\mathbf{I} \ \& \ \mathcal{C}$ | 36 | $84.44 \pm 6.68$ |
| | EasyMKL | $\mathbf{I} \ \& \ \mathcal{C}$ | 36 | $84.78 \pm 6.76$ |
| FW | SimpleMKL | $\mathcal{V} \ \& \ \mathcal{C}$ | 168165 | Out of memory |
| | EasyMKL | $\mathcal{V} \ \& \ \mathcal{C}$ | 168165 | $88.80 \pm 7.02$ |
| FWS | EasyMKLFS | $\mathcal{V} \ \& \ \mathcal{C}$ | 168165 | $96.14 \pm 3.55$ |

Figure B.9 shows the assigned weights of the clinical information by using all the clinical features.

Finally, in Figure B.10 it is possible to note the importance of the clinical data compared to the weight assigned to the voxel of the MRI images.

#### Appendix B.2. Depression

The accuracy results for the Depression dataset are presented in Table B.15.

**Table B.15:** Depression Dataset: comparisons of 5 repetitions of a nested 10-fold crossvalidation balanced accuracy using all the clinical information. The results are divided in 5 families: Baseline, Feature Selection (FS), standard Multiple Kernel Learning (MKL), Feature Weighting by using MKL (FW) and our method in Feature Weighting and Selection (FWS). R corresponds to the number of kernels used.

|  | Algorithm | Kernels | R | Bal. Acc. % |
| --- | --- | --- | --- | --- |
| Baseline | Linear SVM | $\mathbf{C}$ | 1 | $83.33 \pm 15.71$ |
| | SVM | $\mathbf{I} + \mathbf{C}$ | 1 | $67.00 \pm 14.87$ |
| FS | SVM RFE | $\mathcal{V} \ \& \ \mathcal{C}$ | – | $65.24 \pm 11.34$ |
| | SVM t-test | $\mathcal{V} \ \& \ \mathcal{C}$ | – | $63.89 \pm 10.32$ |
| MKL | SimpleMKL | $\mathbf{I} \ \& \ \mathcal{C}$ | 49 | $84.65 \pm 12.88$ |
| | EasyMKL | $\mathbf{I} \ \& \ \mathcal{C}$ | 49 | $84.56 \pm 13.02$ |
| FW | SimpleMKL | $\mathcal{V} \ \& \ \mathcal{C}$ | 713864 | Out of memory |
| | EasyMKL | $\mathcal{V} \ \& \ \mathcal{C}$ | 713864 | $84.55 \pm 12.19$ |
| FWS | EasyMKLFS | $\mathcal{V} \ \& \ \mathcal{C}$ | 713864 | $84.21 \pm 10.72$ |

Figure B.11 and B.12 depict the assigned weights of the clinical information by using all the clinical features and the ration between the weight assigned to the clinical data with respect to the weight assigned to the different fMRIs.

**Figure B.9:**
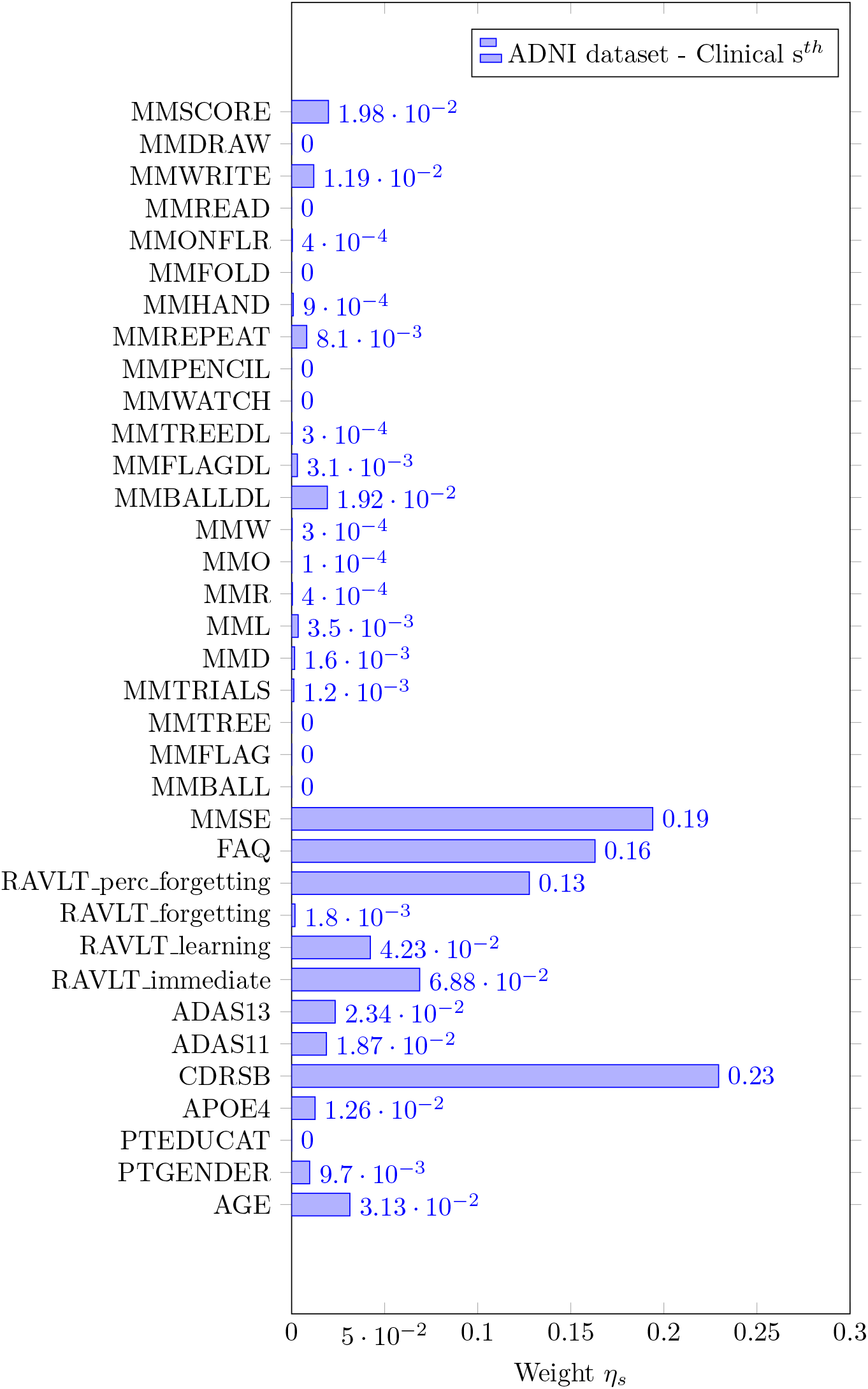
EasyMKL assigned weights for the all the clinical information for the ADNI dataset.

**Figure B.10:**
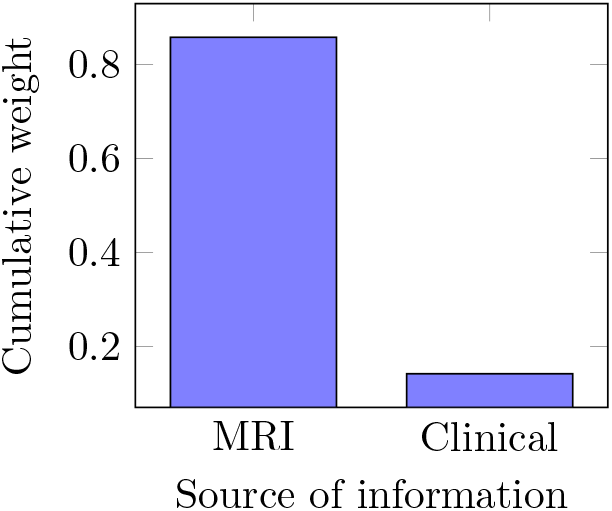
EasyMKLFS assigned weights for the different sources of information: MRI image and all the clinical measurements.

**Figure B.11:**
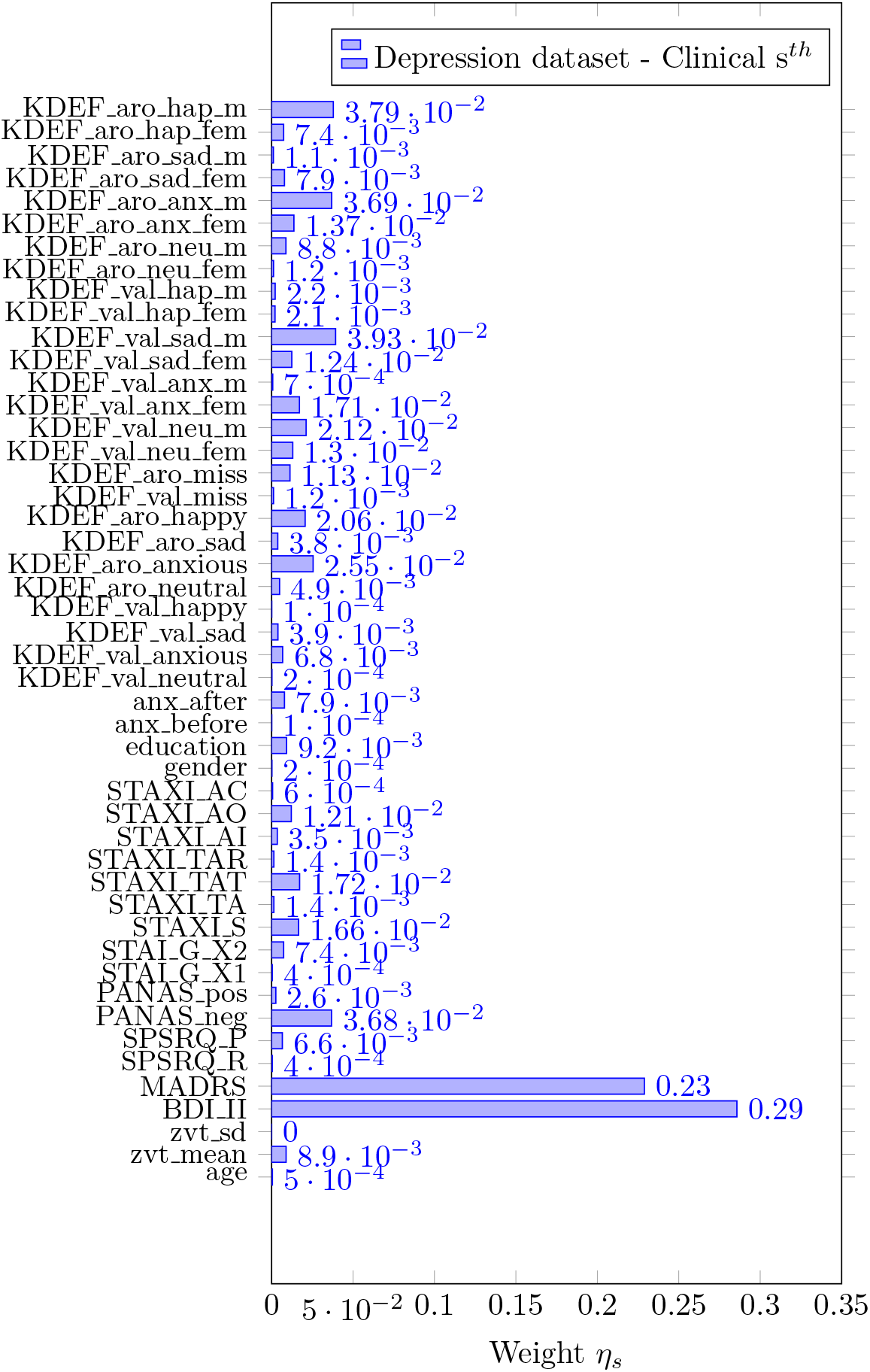
EasyMKL assigned weights for the clinical information for the Depression dataset.

**Figure B.12:**
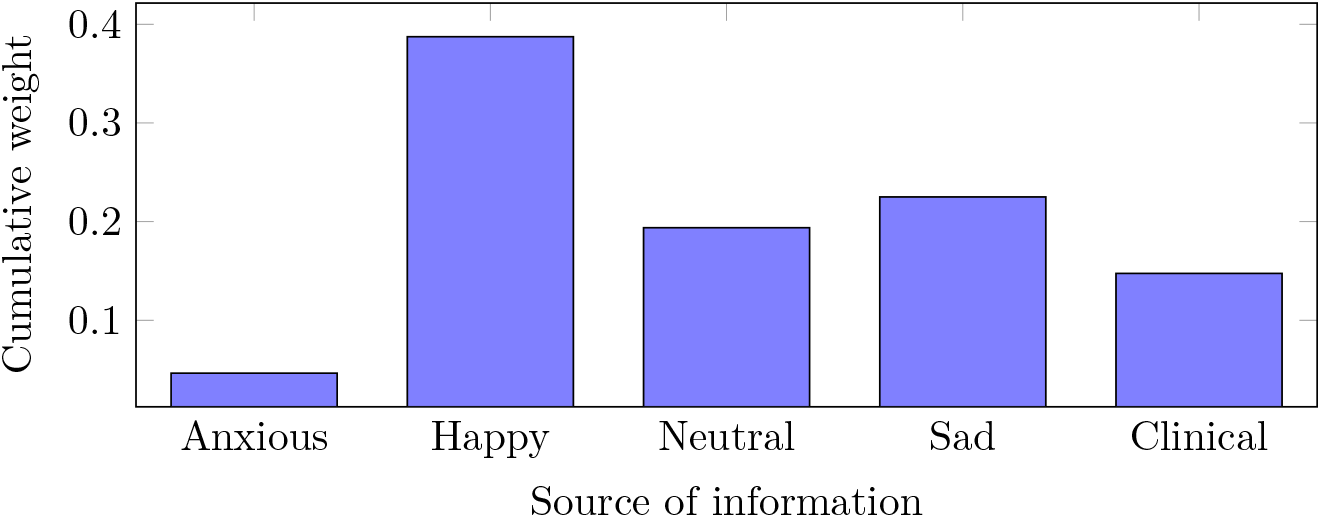
EasyMKLFS assigned weights for the different sources of information of the Depression dataset: Anxious image, Happy image, Neutral image, Sad image and clinical measurements.

## Footnotes

2 A norm is a function that assigns a strictly positive length or size to a vector.

3 www.cse.msu.edu/~cse902/S14/ppt/MKL_Feb2014.pdf

4 EasyMKL implementation: github.com/jmikko/EasyMKL

5 We performed the same experiments as presented in Section 4 using RBF kernels instead of linear ones and we obtained comparable results with an higher computational requirements. For this reason we decided to maintain only the linear kernels in our setting. It is important to note that, in general, our method can be applied to any family of kernels.

